# Pushing the limits of single molecule transcript sequencing to uncover the largest disease-associated transcript isoforms in the human neural retina

**DOI:** 10.1101/2024.09.10.612265

**Authors:** Merel Stemerdink, Tabea Riepe, Nick Zomer, Renee Salz, Michael Kwint, Raoul Timmermans, Barbara Ferrari, Stefano Ferrari, Alfredo Dueñas Rey, Emma Delanote, Suzanne E. de Bruijn, Hannie Kremer, Susanne Roosing, Frauke Coppieters, Alexander Hoischen, Frans P. M. Cremers, Peter A.C. ’t Hoen, Erwin van Wijk, Erik de Vrieze

**Affiliations:** Department of Otorhinolaryngology, Radboud University Medical Center, Nijmegen, 6525 GA, The Netherlands; Donders Institute for Brain, Cognition and Behaviour, Radboud University Medical Center, Nijmegen, 6525 GA, The Netherlands; Department of Medical BioSciences, Radboud University Medical Center, Nijmegen, 6525 GA, The Netherlands; Department of Human Genetics, Radboud University Medical Center, Nijmegen, 6525 GA, The Netherlands; Fondazione Banca degli Occhi del Veneto, Zelarino – Venice, 30174, Italy; Center for Medical Genetics, Ghent University Hospital, Ghent, 9000, Belgium; Department of Biomolecular Medicine, Ghent University, Ghent, 9000, Belgium; Department of Pharmaceutics, Ghent University, Ghent, 9000, Belgium; Department of Internal Medicine and Radboud Center for Infectious Diseases (RCI), Radboud University Medical Center, Nijmegen, 6525 GA, The Netherlands

## Abstract

Sequencing technologies have long limited the comprehensive investigation of large transcripts associated with inherited retinal diseases (IRDs) like Usher syndrome, which involves 11 associated genes with transcripts up to 19.6 kb. To address this, we used PacBio long-read mRNA isoform sequencing (Iso-Seq) following standard library preparation and an optimized workflow to enrich for long transcripts in the human neural retina. While our workflow achieved sequencing of transcripts up to 15 kb, this was insufficient for Usher syndrome-associated genes *USH2A* and *ADGRV1*, with transcripts of 18.9 kb and 19.6 kb, respectively. To overcome this, we employed the Samplix Xdrop System for indirect target enrichment of cDNA, a technique typically used for genomic DNA capture. This method facilitated the successful capture and sequencing of *ADGRV1* transcripts as well as the full-length 18.9 kb *USH2A* transcripts. By combining algorithmic analysis with detailed manual curation of sequenced reads, we identified novel isoforms and alternative splicing events across the 11 Usher syndrome-associated genes, with implications for diagnostics and therapy development. Our findings demonstrate the Xdrop system’s adaptability for cDNA capture and the advantages of integrating computational and manual transcript analyses. The full neural retina sequencing dataset is available via EGA under identifier EGAD50000000720.

## INTRODUCTION

The human retina is a complex multicellular tissue that plays a crucial role in visual function by converting light into the electrical signals that are interpreted by the brain. Understanding the molecular composition of the human retina, particularly the wide variety of transcript isoforms expressed there, is essential for comprehending disease mechanisms and designing effective treatment strategies for inherited retinal diseases (IRDs) (Braun *et al*., 2013). A key factor contributing to the diversity of transcript isoforms expressed in the retina is alternative splicing, a biological process that allows a single gene to encode multiple transcript isoforms leading to tissue-specific differences in gene expression and function. This process involves the use of alternative transcription initiation and termination sites, intron retention, exon skipping, and alternative splice donor and acceptor sites. The retina is a highly specialized tissue that is known to be enriched for these tissue-specific alternative splicing events (Cao *et al*., 2011; Liu and Zack, 2013).While previous studies using RNA short-read sequencing provided valuable insights, they often fall short in fully characterizing the diverse array of transcript isoforms present in the human retina (Ciampi *et al*., 2022; Murphy *et al*., 2016; Ruiz-Ceja *et al*., 2023; Sarantopoulou *et al*., 2021).

The PacBio long-read mRNA isoform sequencing (Iso-Seq) technology (Wang *et al*., 2016) provides deeper insights into the transcriptome complexity caused by alternative splicing events. This technology eliminates the need for *de novo* assembly, and consequently provides greater certainty in the identification of alternative splicing events, as it can generate full-length transcripts up to 10 kb in length. This is significant, as the average human transcript length is approximately 2 kb. However, many transcripts associated with IRDs are considerably longer, and are therefore at (or beyond) the limit of what can be reliably investigated using PacBio Iso-Seq analyses. The longest two annotated transcripts of IRD-associated genes are *USH2A* (18.9 kb) and *ADGRV1* (19.6 kb), both associated with Usher syndrome, which is an autosomal recessively inherited disorder characterized by the combination of sensorineural hearing loss and the progressive loss of visual function due to retinitis pigmentosa (RP). The disorder is clinically and genetically diverse, with 11 associated genes. With the exception of *CIB2* (USH1J), the full-length isoforms of the Usher syndrome-associated genes surpass the average human transcript length of 2 kb, with 5 of them exceeding the mean transcript length of RETNET genes (4.7 kb).

Obtaining a comprehensive understanding of the Usher syndrome-associated transcript isoforms in the human retina is crucial, as it facilitates the development of therapeutic strategies and enables accurate classification of genetic variants linked to the disorder. To this end, we aimed to provide an overview of the Usher syndrome-associated transcript isoforms in the human neural retina, utilizing the PacBio long-read mRNA Iso-Seq dataset from our previous study (Riepe *et al*., 2024), and supplementing it with two additional datasets aimed at capturing full-length reads from even the longest known transcript isoforms. Additionally, we performed an in-depth analysis that specifically focused on the 11 genes associated with Usher syndrome.

The Iso-Seq workflow for sample preparation involves extraction of high-quality RNA, synthesis of cDNA, size-selection and amplification of transcripts, followed by a SMRTbell library preparation and sequencing. Performing a size-selection is crucial to enable sequencing transcripts of above-average lengths, as the PacBio sequencing platform has the tendency to preferentially sequence shorter transcripts. The size-selection step in the standard Pacbio Iso-Seq workflow is optimized for transcripts centered around 2 kb, which corresponds to the average transcript length in human, making it suitable for genome-wide studies as we performed in Riepe *et al*. (2024). However, as nearly all Usher syndrome-associated genes have transcripts exceeding 2 kb, we generated an additional sequencing dataset using an optimized long transcript workflow to enable the sequencing of larger transcripts up to > 10 kb. Although this workflow was better optimized to enable sequencing full-length transcripts of Usher syndrome-associated genes *MYO7A* (7.4 kb), *CDH23* (11.1 kb), *PCDH15* (9.4 kb) and *WHRN* (4.1 kb), this remained insufficient for sequencing the longest *USH2A* (18.9 kb) and *ADGRV1* (19.6 kb) transcript isoforms. In an attempt to further enhance the capture of full-length transcripts for these two genes, we also employed an ‘indirect targeted enrichment’ approach using the Samplix Xdrop System (Madsen *et al*., 2020), followed by PacBio long-read sequencing. By integrating the data from these three sequencing workflows and employing a combined strategy of algorithmic analysis and manual curation, we were able to obtain a comprehensive overview of the Usher syndrome-associated transcript isoforms present in the human neural retina. The approach demonstrated here illustrates the advantages of integrating algorithmic and extensive manual analyses to fully capture the transcriptomic complexity within tissues such as the human neural retina. Therefore, the analysis applied here represents a valuable approach that can be applied to all IRD-associated genes to advance diagnostic pipelines and the development of therapeutic strategies.

## MATERIAL AND METHODS

### Tissue collection for PacBio Iso-Seq long-read mRNA sequencing

Human donor eyes, deemed unsuitable for corneal transplantation, were used in this study. Written consent was obtained from the donors’ next of kin in accordance with the guidelines of the Italian National Transplant Centre (Centro Nazionale Trapianti, Rome, Italy). The procurement and processing of these tissues followed Italian law and adhered to the principles of the Declaration of Helsinki and the guidelines of the European Eye Bank Association. Three human neural retinal samples were obtained from non-visually impaired individuals through the Fondazione Banca degli Occhi del Veneto (Venice, Italy). Prior to this study, we performed whole-genome sequencing on DNA isolated from these retinal samples to screen for and exclude any known genetic variants associated with inherited retinal disorders. The eyes were enucleated within 2-12 hours postmortem, and the retinal extraction followed established protocols (Niyadurupola *et al*., 2011; Osborne *et al*., 2016). After cornea removal, the eyeball was dissected at the *ora serrata*; iris, lens, and vitreous body were removed before carefully detaching the retina from sclera and retinal pigment epithelium by cutting at the optic nerve head. Subsequently, retinal samples were snap-frozen in cryovials using liquid nitrogen and shipped on dry ice. Detailed information about the donors is presented in Table S1.

### RNA isolation and PacBio Iso-Seq library preparation following the standard workflow

Total RNA was extracted from the human neural retina samples by adding 500 µL of Trizol reagent to each sample, which was then homogenized in a 2 mL tube containing a sterile glass bead using a TissueLyser (QIAGEN, Aarhus, Denmark) in two cycles at 30 Hz for 30 seconds. Following a 5-minute incubation at room temperature (RT), 100 µL chloroform was added to the samples which were then mixed and incubated for another 3 minutes at RT before being centrifuged at 12,000 g for 15 minutes. The resulting aqueous phase was collected and mixed with 1 µL glycogen (5 µg/µl) and an equal volume of isopropanol. This mixture was incubated at 20°C for 75 minutes followed by centrifugation at 12,000 g for 30 minutes at 4°C. Subsequently, the supernatant was removed and the remaining RNA pellet was dissolved in MQ water and further purified and DNAse treated using the Nucleospin RNA Clean-up Kit (Macherey-Nagel, Düren, Germany) according to the manufacturer’s instructions. The total isolated RNA quantity was measured with a Qubit fluorometer (Thermo Fisher Scientific, Waltham, MA, USA). Additionally, RNA integrity number (RIN) values were determined using a 2100 Bioanalyzer (Aligent Technologies, Santa Clara, CA, USA). The RIN values of all three samples exceeded 7.0 (Table S1), and 300 ng of RNA input was utilized to generate Iso-Seq SMRTbell libraries following the Iso-Seq-Express-Template-Preparation protocol version 2.0 (Pacific Biosciences, California, USA).

These libraries were prepared using the standard workflow, suitable for samples with a predominant transcript size of approximately 2 kb, and the SMRTbell® library binding kit 2.1 (Pacific Biosciences, California, USA). The samples were not labeled with a barcode, and 500 ng of cDNA was used for the subsequent steps in the procedure. The quantification and assessment of the SMRTbell library for each sample were conducted using Qubit (Thermo Fisher Scientific, Waltham, MA, USA) and an Agilent Bioanalyzer 2100 employing HS RNA screentape. The on-plate loading concentration of the final Iso-Seq SMRTbell libraries was set at 80 pM, and sequencing was carried out on a Sequel IIe system (Pacific Biosciences, California, USA) with a movie time of 24 hours.

### PacBio Iso-Seq library preparation following adjusted workflow optimized for large transcripts

In addition to the three libraries prepared following the PacBio standard workflow for samples with a predominant transcript size of approximately 2kb, an additional library was prepared and optimized to obtain amplified cDNA enriched for larger transcripts. 300 ng of RNA sample 2 was used to generate the Iso-Seq SMRTbell library using the Iso-Seq-Express-Template-Preparation protocol version 2.0 (Pacific Biosciences, California, USA). The library was prepared using the PacBio long transcript workflow suitable for samples with a transcript size larger than 3 kb utilizing the SMRTbell® library binding kit 2.1. To further enhance the enrichment for large transcript sizes, deviations in the cDNA amplification and purification process involved carrying out 5 additional PCR cycles after the ‘Purification of Amplified cDNA’ step in Appendix 1 of the protocol. After the additional amplification, a size-selection step was conducted using a 3.3X ratio of diluted AMPure PB beads according to the PacBio Procedure Using AMPure PB Beads for Size-Selection protocol 101-854-900 version 2.0 (Pacific Biosciences, California, USA). The concentration and size distribution of the resulting SMRTbell library was evaluated using Qubit (Thermo Fisher Scientific, Waltham, MA, USA) and an Agilent Bioanalyzer 2100 with HS RNA screentape. The on-plate loading concentration of the final Iso-Seq SMRTbell libraries was set at 120 pM, and sequencing was carried out on a Sequel IIe system (Pacific Biosciences, California, USA) with a movie time of 30 hours.

### PacBio long-read Iso-Seq data analysis

Four PacBio long read-RNA Iso-Seq samples (three samples prepared following the PacBio standard workflow, 1 sample prepared following the optimized PacBio long-transcript workflow) were analyzed with IsoQuant (Prjibelski *et al*., 2023). Three separate datasets were created: the first dataset included the IsoQuant analysis of the three standard workflow samples, like we previously performed in Riepe *et al*. (2024). The second dataset combined the analysis of PacBio standard workflow retina samples 1, 2, and 3 with the retina sample prepared following the optimized long transcript workflow. Lastly, the third dataset consists only of the long-transcript workflow sample, analyzed with less strict filtering settings. The sequencing data resulting from the PacBio Iso-Seq approaches, that were used for IsoQuant analyses are available via EGA with identifier EGAD50000000720.

#### Dataset 1

We generated circular consensus sequence (CCS) reads with a minimum accuracy of 0.99, maximum length of 30,000, and minimum number of three passes using CCS (v6.2.0). Primers and SMRT adapters were removed with Lima (v2.4.0) to generate full-length (FL) reads. Iso-Seq3 (v.3.4.0) refine was applied to obtain full-length non concatemer (FLNC) reads with parameter --require-polya. FLNC reads were converted from BAM to FASTQ and then aligned using Minimap2 (v2.24) (Li, 2021) with parameters -ax splice -uf --secondary=no -C5 -O6,24 –B4 -MD. The mapped isoforms were classified with IsoQuant using the GENCODE v39 primary assembly annotation and parameters -- *complete_gened, --datatype pacbio_ccs, --fl_data, --sqanti_output, --count_exons, -- model_construction_strategy fl_pacbio, --check_canonical, --transcript_quantification unique_only, -- gene_quantification unique_only, and --splice_correction_strategy default_pacbio.* Additionally, we created a separate dataset for which the reads were clustered with Iso-Seq3 cluster with parameter --useqvs before alignment for manual assessment in IGV. Minimap2 was run with the same parameters, excluding the -MD flag.

#### Dataset 2

This dataset consists of the PacBio standard workflow samples and the PacBio long-transcript workflow sample. The analysis was performed similarly to dataset 1.

#### Dataset 3

This dataset consists only of the PacBio long-read RNA sequencing sample enriched for long transcript sizes. The analysis was performed similarly to dataset 1, however, we used less strict filtering criteria. For CCS, we changed the minimum accuracy to 0.80 and the minimum number of passes to 0. We then continued the analysis as described for dataset 1.

### Data analysis and transcript visualization

RStudio software (v4.3.2) was used to obtain read and transcript counts from the IsoQuant output files. GGtranscript package (Gustavsson *et al*., 2022) was used to visualize transcripts from the GTF output files. ORFs were predicted with Snapgene. Details and R scripts are posted on GitHub (https://github.com/erikdevrieze/USH_retina).

### Targeted Iso-Seq for *USH2A* and *ADGRV1* transcripts using the Samplix Xdrop Sort

Next to the conventional Iso-Seq transcriptome analysis, the Samplix Xdrop Sort (Samplix ApS, Birkerød, DK) was employed for a targeted Iso-Seq analysis, enriched for *USH2A* and *ADGRV1* transcripts. Of each retina sample, 300ng of RNA was utilized for cDNA synthesis using the NEBNext Single Cell/Low Input cDNA Synthesis kit (New England Biolabs, Ipswich, MA, USA). For this purpose, the “Preparing Iso-Seq libraries using SMRTbell prep kit 3.0” protocol version 2.0 (Pacific Biosciences, California, USA) was followed until the cDNA amplification step. The synthesized cDNA of all three samples was pooled and the final DNA concentration of the resulting pool was determined using the Qubit with the single stranded DNA kit (Thermo Fisher Scientific, Waltham, MA, USA) according to the standard workflow. In order to perform the targeted enrichment, single cDNA molecules were packaged in double emulsion (DE) droplets together with detection sequence primers using the Samplix Xdrop Sort. The detection sequence primers were designed to target a 100-150bp region located in either the 5’, middle or 3’ region of full-length *USH2A* (ENST00000307340.8) and *ADGRV1* (ENST00000405460.9) transcripts respectively, as to increase the likelihood of capturing full-length transcripts as well as known and novel shorter isoforms. Details of the detection sequence primers, or droplet PCR (dPCR) primers, and the exons they target can be found in Table S2.

The encapsulation process was performed separately for each of the six dPCR primer sets to ensure targeting of a single region during enrichment. Samples were run in technical duplicates, and two positive controls for amplification and sorting provided by Samplix were included on each Xdrop DE20 Cartridge (Samplix ApS, Birkerød, Denmark). The input concentration was 2.8 ng of pooled cDNA. The sample mix, prepared using the “Setup of droplet PCR” and “Positive control DNA enrichment droplet PCR” sections of the Samplix “Targeted enrichment of DNA with Xdrop or Xdrop Sort” protocol version 2.0, was loaded onto the Xdrop DE20 cartridge. Droplets were produced according to the “Producing DE20 droplets” guide, and the droplet PCR was initiated as described in the “DE droplet production” and “Droplet PCR reaction” sections of the protocol.

During PCR amplification, droplets containing cDNA molecules with the target sequence produced fluorescent signals detectable by the Xdrop Sort. Sorting followed the “DE20 droplet sorting with Xdrop Sort” quick guide, with samples incubated for 15 minutes at room temperature after adding the staining buffer. Positive controls were included in each sorting run, and after sorting, transcripts were released from droplets following the “Breaking DE20 droplets containing DNA” guide.

Since the cDNA amount after sorting was insufficient for direct sequencing, the captured transcripts were amplified using multiple displacement amplification (MDA) with non-sequence-specific primers to avoid bias towards smaller transcripts. Single transcripts were packaged into SE droplets using the “Producing SE85 droplets” guide and the Samplix Xdrop Sort Manual version 4.0, with 10 µl of enriched cDNA as input. Additionally, 1 pg of unenriched cDNA and a non-template control were included. SE droplets were incubated at 30°C for 16 hours, 65°C for 10 minutes, and 4°C until DNA release. The “Breaking SE85 droplets” guide was followed for droplet breaking.

Enrichment validation was performed using qPCR to compare enriched samples with the original, un-enriched cDNA pool. The amplified 1 pg of unenriched cDNA and non-template control from the MDA served as positive and negative controls, respectively. qPCR primers, designed to target 100-125 bp regions close to the targeted regions of the dPCR primers, were used in technical replicates to negate bias from unintended target capture. Primer compositions and targeted exons are detailed in Table S2. Each qPCR reaction contained 1 µl of template cDNA, 10 µl of Gotaq 2x Master Mix (Promega Corporation, Madison, USA), 1 µl of forward and reverse qPCR primers, and 7 µl of nuclease-free water. Reactions were run on a QuantStudio 3 Real-Time PCR System (Thermo Fisher Scientific, Waltham, MA, USA) with the following program: 50°C for 1 second, 95°C for 10 minutes, 40 cycles of 95°C for 15 seconds and 60°C for 30 seconds, then 95°C for 15 seconds, 60°C for 1 minute, and finally 95°C for 15 seconds.

### PacBio library preparation and sequencing - adjusted workflow for Samplix enriched transcripts

To remove the branched structures present in the amplified transcripts generated through the MDA process, an enzymatic digestion was performed following the “Sequencing library prep recommendations for sequencing after the Samplix targeted enrichment workflow” protocol version 3.1 (Samplix ApS, Birkerød, Denmark). A total of 500 ng of MDA-amplified and purified cDNA was used to create multiplexed SMRTbell amplicon libraries, following the manufacturer’s instructions for the SMRTbell prep kit 3.0 (Pacific Biosciences, California, USA). If the sample had less than 500 ng of cDNA, linearized plasmids not containing the region of interest were added to spike the samples. The samples were then prepared for sequencing using the Sequel IIe Binding Kit 3.2 (Pacific Biosciences, California, USA), following the recommended protocols. Finally, 115 µl of the final mix was loaded per well and long read sequencing was performed using the Sequel IIe system (Pacific Biosciences, California, USA). Following sequencing, the subreads were demultiplexed using lima V.2.5.0, and combined to create a consensus sequence using CCS V.6.3.0. These reads were filtered RQ 0.99 to obtain HiFi reads, and HiFi reads were then mapped along the GRCh38 reference genome using pbmm2 V1.8.0 with the –preset isoseq mode.

### Tissue collection for Oxford Nanopore Technology (ONT) sequencing

For the independent ONT validation dataset, rest material from human donors (n=3) without any known or clinical evidence of retinal disease was collected from the tissue banks of either Ghent University Hospital or Antwerp University Hospital. This collection adhered to the ethical standards of the Declaration of Helsinki and received approval from the Ethics Committee of Ghent University Hospital (IRB approval B670201837286). Details about the donors can be found in Table S3. The eyes were transported in CO2 Independent Medium (Gibco) prior to dissection. To preserve RNA integrity and minimize the effects of autolysis, retinas were only harvested from eyes with a total post-mortem interval of less than 20 hours. Following a visual inspection to ensure no contamination from the retinal pigment epithelium (RPE), the neural retinas were either processed immediately for total RNA isolation or snap-frozen and stored at −80°C for later use.

### RNA isolation and Oxford Nanopore Technology (ONT) sequencing

Total RNA was extracted from the post-mortem adult human neural retina samples using the RNeasy Mini kit® (Qiagen), following the manufacturer’s instructions. The extracted RNA then underwent DNase treatment (ArcticZymes, Tromsø, Norway) followed by poly-A capture. Samples of poly-A mRNA with sufficient quality (RNA Integrity Value, RIN > 8.0) were used for direct-cDNA library preparation using the SQK-DCS109 kit (Oxford Nanopore Technologies (ONT)), with minor modifications to the supplier’s protocol. Each prepared library was then loaded onto a FLO-PRO002 flow cell (ONT) and sequenced on an ONT PromethION device for 72 hours. Details regarding the number of reads can be found in Table S3.

### ONT data analysis of selected genes

MinKNOW version 5.1.0 was used to produce fast5 files, which were subsequently base-called with Guppy version 6.1.5. The reads were then aligned to the Human Reference Genome GRCh38/hg38 using minimap2 version 2.24 with the -ax splice flags for spliced alignments. Alignment files were converted to BAM format, sorted, and indexed using SAMtools version 1.15 to produce the dataset investigated here.

### Relative expression of *MYO7A* isoforms

Quantitative PCR analysis was conducted to determine the relative expression levels of the different *MYO7A* transcript isoforms identified in the PacBio Iso-Seq datasets across three human neural retina samples. The same RNA used for preparing the PacBio libraries was employed as the template, with 250 ng of RNA being used for cDNA synthesis using the SuperScript IV Reverse Transcriptase kit (Thermo Fisher Scientific, Waltham, MA, USA, #18090200). Quantitative PCR analysis was performed using GoTaq qPCR Master Mix (Promega), following the manufacturer’s protocol. Transcript-specific primers were designed and validated to target the 5’ canonical start site, the 5’ alternative start site, and the 3’ end of the identified *MYO7A* isoforms, as well as the reference gene *GUSB*. Amplifications were carried out using the QuantStudio 3 Real-Time PCR system (Applied Biosystems, Waltham, MA, USA), with PCR reactions performed in triplicate. Relative gene expression levels compared to the reference gene *GUSB* were determined using the 2−ΔCt method. The primers used are listed in Table S4.

### Analysis of protein isoforms

To predict the 2D protein domain architecture of encoded proteins in silico analyses were performed using the SMART online tool (Letunic *et al*., 2021). 3D protein structures were modelled using AlphaFold (Jumper *et al*., 2021) using the Google Colab notebook (v1.5.3) with standard settings. The 3D structures were visualized with YASARA (Krieger and Vriend, 2014), and structural alignments were conducted using the MUSTANG algorithm (Konagurthu *et al*., 2006).

## RESULTS

High-quality RNA was extracted from three human neural retina samples to enable PacBio long-read mRNA Iso-Seq, to gain an overview of the Usher syndrome-associated transcript isoforms expressed in the human retina. We supplemented our existing sequencing dataset obtained with the standard workflow (published in Riepe *et al*. (2024)), with a dataset obtained from an optimized PacBio long transcript workflow, enabling the sequencing of larger transcripts up to 15 kb. Additionally, to capture and sequence the largest transcripts of the *USH2A* (18.9 kb) and *ADGRV1* (19.6 kb) genes, we employed an indirect targeted enrichment approach using the Samplix Xdrop System, followed by PacBio long-read sequencing. By integrating the data from these three sequencing workflows and combining algorithmic analysis with manual curation (Figure 1), we obtained a comprehensive overview of the landscape of Usher syndrome-associated transcript isoforms in the human neural retina. Finally, selected transcripts and events were validated using an independent Oxford Nanopore Technology (ONT) long-read sequencing retina dataset.

**Figure 1:**
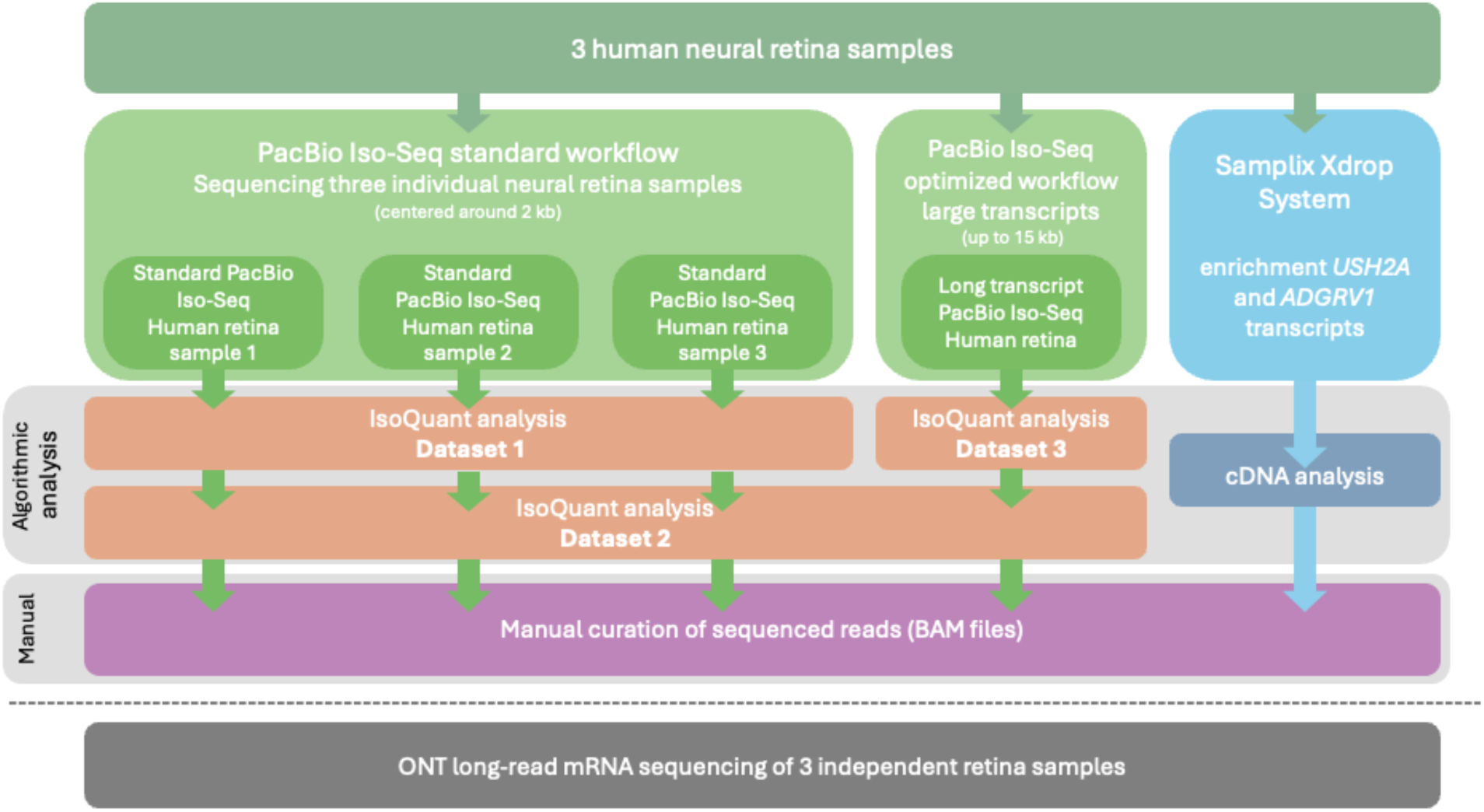
Overview of the Sequencing Workflows and Subsequent Analyses. The figure illustrates the sequencing workflows and subsequent analysis performed on RNA extracted from three human neural retina samples. The workflows included PacBio long-read mRNA Iso-Seq using both the standard and an optimized long transcript workflow. The analysis was carried out in three distinct datasets: Dataset 1 comprised the standard workflow samples analyzed with IsoQuant, Dataset 2 involved a combined analysis of the standard and optimized long transcript workflows, and Dataset 3 focused solely on data from the long transcript workflow. Additionally, an indirect targeted enrichment of transcripts for the *USH2A* and *ADGRV1* genes was achieved using the Samplix Xdrop System, followed by PacBio long-read sequencing and cDNA analysis. Additionally, the sequenced reads were manually curated using BAM files in the Integrative Genomics Viewer. Integrating data from the three sequencing workflows and combining algorithmic analysis with manual curation enabled a thorough characterization of the Usher syndrome-associated transcript isoforms present in the human neural retina. Validation of selected transcripts and events was performed using an independent Oxford Nanopore Technology (ONT) long-read sequencing dataset of three independent retina samples.

Table 1 summarizes the known Usher syndrome-associated retinal transcripts identified through IsoQuant analysis (Prjibelski *et al*., 2023) of the PacBio standard workflow and the optimized long transcript workflow. Additionally, it highlights frequently observed novel transcript isoforms and events that have not been previously annotated, which were confirmed by Oxford Nanopore Technology (ONT) long-read mRNA sequencing of independent retina samples. Furthermore, a detailed overview of minor events observed following the manual curation of sequenced reads using the BAM files in the Integrative Genomics Viewer (IGV) is presented in Table S5. The findings for *MYO7A* (Figure 3), *WHRN* (Figure 4), *USH2A* (Figure 5 and 6) and *ADGRV1* (Figure 7 and 8) are showcased to illustrate the different types of observations in the dataset, such as the identification of novel, previously unidentified isoforms, novel coding exons, or regions sensitive to pseudoexon (PE) inclusions. For the remaining Usher syndrome-associated genes, an overview of the algorithmic output and manual curation results has been generated and presented in Figures S3-S9.

**TABLE 1:**
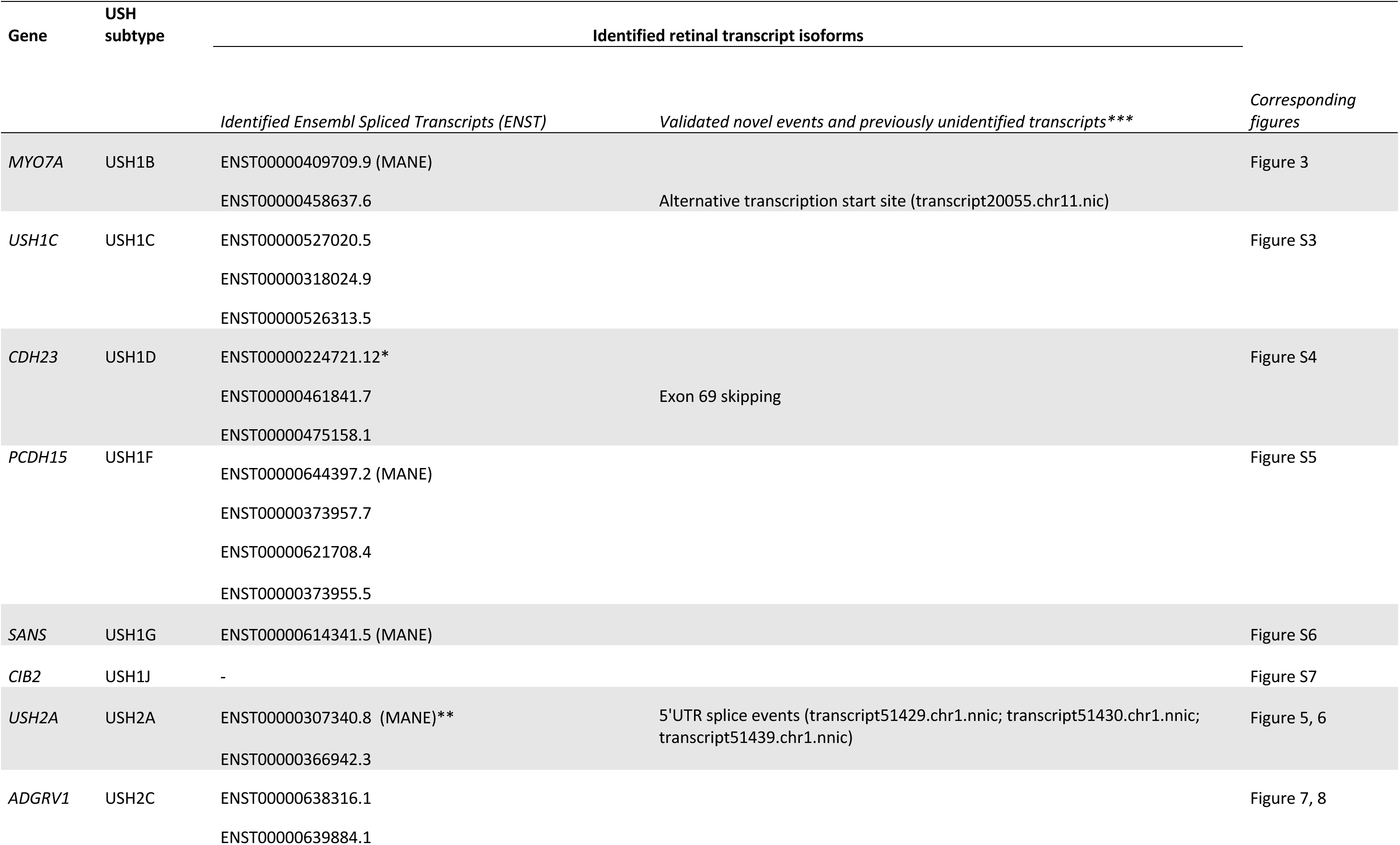
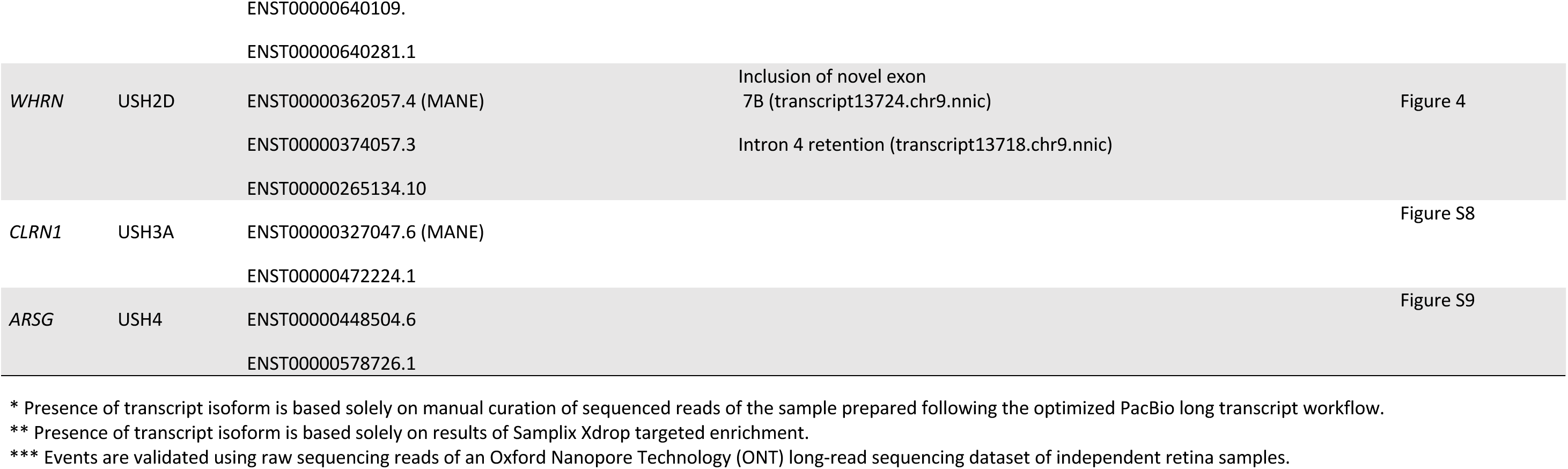
Identified USH-associated retinal transcript isoforms and prevalent events observed across the majority of transcripts.

### Exploring Usher syndrome-associated transcript isoforms through Iso-Seq PacBio standard- and optimized long transcript workflows

After quality control, PacBio transcripts were analyzed and classified using the IsoQuant classification (Prjibelski *et al*., 2023). Transcripts identified from the PacBio standard workflow had an average read length of 2.6 kb, which is shorter than most Usher syndrome-associated genes. In contrast, the optimized PacBio long transcript workflow successfully enriched for larger transcript sizes, producing transcripts up to 15 kb with an average size of 4.7 kb (Figure 2A). Transcripts for all 11 Usher syndrome-associated genes were identified in the samples prepared according both the PacBio standard and long transcript workflow, except for the smallest Usher gene, *CIB2* (1.5 kb), which was detected in retina sample 2 and 3, but not in retina sample 1 (Figure S1). The optimized long transcript workflow increased the number of reads for Usher genes with coding sequences larger than the average GENCODE read length of 2.4 kb (Figure 2C).

**Figure 2:**
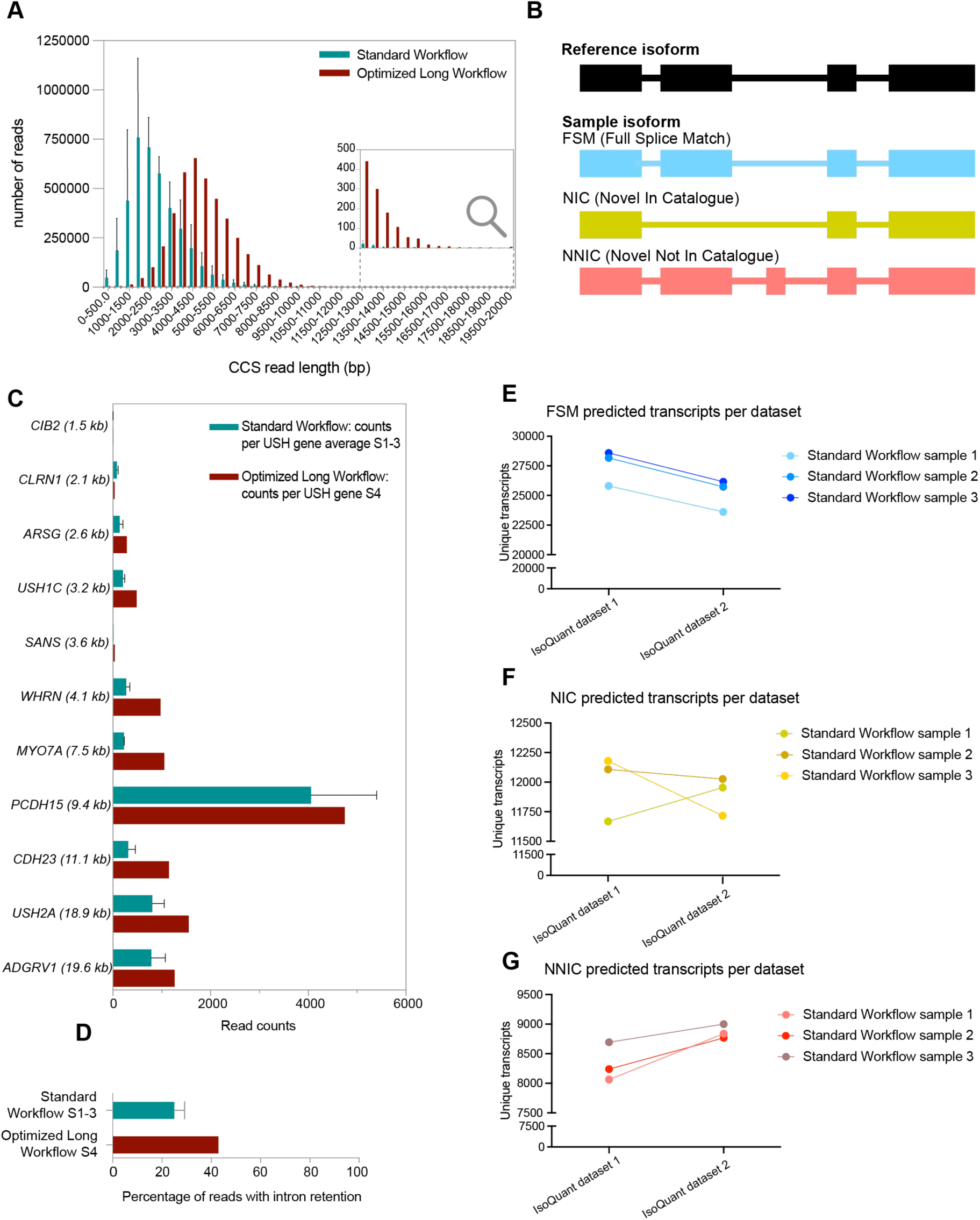
Exploring the Usher syndrome-associated transcript isoform landscape in the human neural retina using PacBio long-read mRNA Iso-Seq A. The size distribution of sequenced transcripts derived from the standard workflow (blue) and optimized long workflow (red) datasets. For the standard workflow dataset, the mean size distribution across the three sequenced samples is depicted ± standard deviation (SD). **B.** Schematic representation that compares sample transcripts to the GENCODE reference transcriptome. Transcript isoforms that fully match the reference are classified as Full Splice Matches (FSMs). Novel transcripts are those that either align to the reference splice junctions, in which case they are designated as Novel In Catalog (NIC), or contain novel splice junctions and are termed Novel Not In Catalog (NNIC). **C.** Comparison of Usher syndrome-associated transcript coverage between the standard workflow and optimized long workflow dataset. The Usher genes are arranged in order from smallest to largest coding sequence, with the coding sequence length of the largest known transcript for each gene provided in brackets. For the standard workflow dataset, the mean +/− SD transcript length across the three sequenced samples is presented **D.** Quantification of the percentage of reads displaying intron retention in standard workflow samples 1-3 (mean of 3 samples ± SD) versus long workflow sample 4. **E, F, G.** Number of unique transcripts associated with each transcript class, as predicted by IsoQuant for each standard workflow sample. Based on the input per dataset, classifications differ for IsoQuant dataset 1 (analysis of 3 Standard Workflow samples) or IsoQuant dataset 2 (combined analysis of 3 standard workflow samples with the optimized long transcript workflow sample). **E.** Number of unique FSM predicted transcripts per dataset, **F.** Number of unique NIC predicted transcripts per dataset, **G.** Number of unique NNIC predicted transcripts per dataset.

In Riepe *et al*. (2024), we integrated PacBio long-read mRNA Iso-Seq data with proteomics and genomic data to construct a proteogenomic atlas. In that publication, the IsoQuant algorithm was utilized to analyze and categorize (Figure 2B) the PacBio Iso-Seq standard workflow samples, resulting in IsoQuant dataset 1. For the current study, we sequenced an additional sample prepared using our optimized PacBio long transcript workflow. Consequently, a second IsoQuant analysis was conducted, combining the sequenced reads from the PacBio standard workflow retina samples 1, 2, and 3 with the retina sample prepared following the optimized long transcript workflow, resulting in the IsoQuant dataset 2 (Figure 1). We noticed, however, that the incorporation of the PacBio long transcript workflow sample in the combined IsoQuant analysis impacted the IsoQuant algorithm’s annotation of Full Splice Matches (FSMs). This becomes evident when comparing the transcript classifications assigned to reads from standard workflow samples 1-3 in our IsoQuant dataset 2 to those in IsoQuant dataset 1. Even though both analyses used the exact same IsoQuant settings, the inclusion of the PacBio long transcript workflow sample in dataset 2 resulted in fewer predicted FSMs for standard workflow samples 1-3 compared to dataset 1, for which the analysis consisted solely of the standard workflow samples 1-3 (Figure 2E). Instead, a shift in the number of Novel In Catalogue (NIC) (Figure 2F) and an increase in the number of Novel Not In Catalogue (NNIC) (Figure 2G) transcripts was observed. This suggests that the incorporation of sequencing reads obtained with the PacBio long transcript workflow influenced the IsoQuant algorithm’s classification of transcript isoforms. The discrepancy in IsoQuant algorithm outputs between the datasets is further demonstrated in Figure S2, which uses the *MYO7A* gene as an illustrative example of the divergence in predicted isoforms by the IsoQuant algorithm when the PacBio long transcript workflow sample is included or excluded.

The difference in the predicted transcript isoforms may be attributed to the disproportionate prevalence of intron-retaining transcripts in the sample prepared following the long transcript workflow. This observation was made through manual inspection of the sequenced reads using the Integrative Genomics Viewer (IGV), which revealed an increased number of intron-retaining transcripts in the sample generated via the long transcript workflow. Quantification indeed revealed a greater percentage of reads displaying intron retention for this sample, compared to the reads from samples 1-3 prepared following the standard workflow (Figure 2D). While the enrichment for larger transcripts successfully enhanced the sequencing coverage of the larger Usher genes, it may also have inadvertently led to an overrepresentation of partially spliced transcripts.

We expected that the IsoQuant algorithm would generate more accurate isoform predictions when analyzing the standard workflow samples independently, rather than analyzing the standard and long workflow samples together. Consequently, for the detailed examination of Usher syndrome-associated transcript isoforms, the study relied on the IsoQuant predictions from dataset 1 and a third dataset was generated in which the long transcript workflow sample was analyzed separately using IsoQuant.

### Iso-Seq identifies a novel predominant *MYO7A* transcript isoform with an alternative 5’ transcription start site in retina

Biallelic variants in *MYO7A* can lead to either non-syndromic hearing loss (DFNB2) (Guilford *et al*., 1994) or, in syndromic cases, cause Usher syndrome type 1B (Weil *et al*., 1995), which represents the most prevalent subtype of Usher syndrome type 1. The *MYO7A* gene encodes the unconventional myosin VIIa motor protein, which is expressed in the outer hair cells of the cochlea (Hasson *et al*., 1995), the retinal pigmented epithelium (RPE), and in the photoreceptor cell connecting cilium (Liu *et al*., 1997; Udovichenko *et al*., 2002). Recent literature describes two major human *MYO7A* transcript isoforms that are expressed in the RPE and the neural retina (Figure 3A, 3B), which differ in the length of exon 35, thereby affecting the structure of the 1^st^ FERM domain in the MYO7A tail (Gilmore *et al*., 2023). IsoQuant analysis confirmed the presence of both isoforms in the human retina (Figure 3A). We observed that the transcript isoform harboring the short form of exon 35 was the most abundant in the retina (4.81 ± 4.04 TPM for ENST00000458637.6 vs. 0.88 ± 0.86 TPM for ENST00000409709.9) (Figure 3C), consistent with the findings of Gilmore *et al*. (2023).

**Figure 3:**
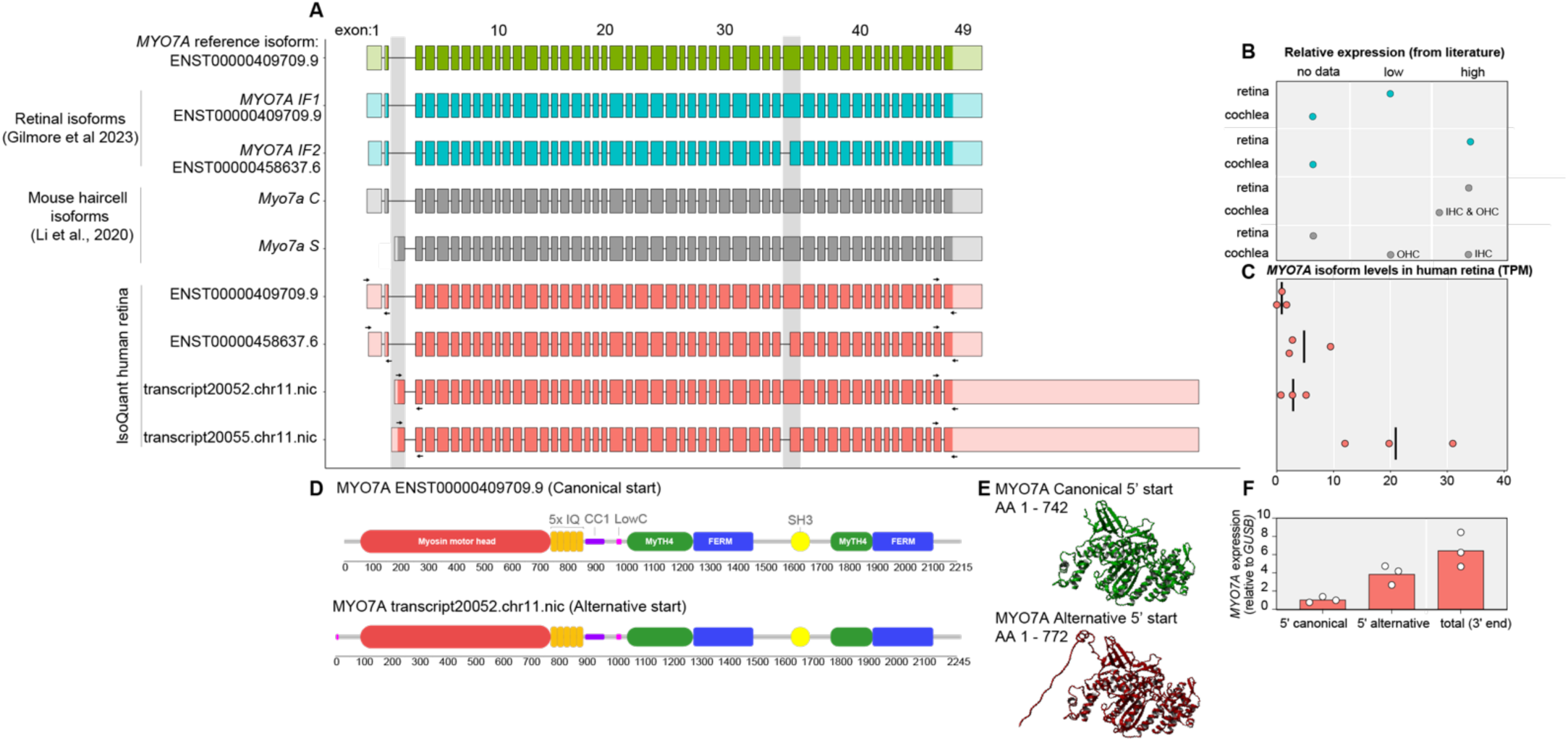
*MYO7A* transcripts identified by IsoQuant analysis compared to known isoforms from the literature. **A.** The GENCODE reference transcript is depicted at the top in green, followed by the known human *MYO7A* transcript isoforms in blue (Gilmore *et al*., 2023) and the murine isoforms in grey (Li *et al*., 2020). The *MYO7A* IsoQuant transcripts are depicted in red. The light green, blue, grey and red colors indicate the untranslated regions (UTR) and the dark green, blue, grey and red colors indicate the open reading frame (ORF) of each transcript. Differences between the IsoQuant transcript isoforms and the GENCODE reference transcript are highlighted in grey boxes. **B.** Relative expression of *MYO7A* isoforms based on literature in either the regna or the cochlea. **C.** The Transcripts Per Million (based on dataset 1) for each IsoQuant isoform are presented for the three individual samples. **D.** The predicted 2D protein domain architecture of the MYO7A protein isoforms with the canonical 5’ start and the alternative 5’ start from transcript20052.chr11.nic. The bar below the 2D protein structures displays the amino acid positions. IQ = isoleucine-glutamine motif, CC1 = Coiled Coil domain, LowC = Low complexity region, MyTH4 = Myosin Tail Homology 4, SH3 = SRC Homology 3 domain. **E.** AlphaFold2 3D protein predictions of the MYO7A protein isoforms, modeled from the 5’ start to the end of the Myosin motor head domain. **F.** RT-qPCR analysis of the relative expression of the *MYO7A* canonical 5’ start site, the alternative 5’ start, and the 3’ end is shown. The locations of the primers for this RT-qPCR are indicated with the arrows on top of the IsoQuant isoforms in Figure 3A.

IsoQuant analysis of dataset 1 identified novel transcript isoforms (transcript20052.chr11.nic; transript20055.chr11.nic) characterized by an alternative 5’ transcription start site coupled with an extended 3’-untranslated region (UTR), which have not been previously reported. Interestingly, a transcript isoform with a similar 5’ start site has been identified in mice (Li *et al*., 2020). Manual curation of sequenced transcripts using the BAM-files in IGV revealed substantial read support for this alternative start site across both the standard- and long transcript workflow samples. Moreover, the use of this alternative transcription start site was also confirmed by ONT long-read mRNA sequencing on three independent retina samples. Protein domain analysis using SMART (Letunic *et al*., 2021) (Figure 3D) and structural modeling with AlphaFold2 (Jumper *et al*., 2021) (Figure 3E) revealed differences between the canonical MYO7A isoform (ENST00000409709.9) and the protein isoform encoded by the IsoQuant transcript with an alternative 5’ start site (transcript20052.chr11.nic). The canonical isoform encodes a protein of 2215 amino acids, whereas transcript20052.chr11.nic is predicted to encode a protein of 2245 amino acids, with a larger N-terminal tail and a predicted low complexity region. To quantify the relative expression of the transcripts with an alternative 5’ start (transcript20052.chr11.nic; transript20055.chr11.nic) compared to the isoforms with the canonical 5’ start, we performed a qPCR using primers specifically targeting the alternative 5’ start, canonical 5’ start, and canonical 3’ end. The results suggest that the identified transcripts with the alternative 5’ start site were the most abundantly expressed transcripts in the human neural retina (Figure 3F).

Further manual curation of sequenced reads using the BAM files in IGV provided an overview of additional splicing events. This analysis uncovered the incidental skipping or inclusion of certain novel exons, which are summarized in Table S5. It is noteworthy that these splicing events are incidental occurrences, except for previously reported 5’ truncation of exon 35 and the observed retention of introns 30 and 37, which were present in approximately 25% of the sequenced transcripts.

### Iso-Seq reveals a novel coding exon and *WHRN* retina specific transcript isoforms

The *WHRN* gene encodes the whirlin (WHRN) protein, and variants in this gene can cause either non-syndromic recessive deafness (DFNB31) or Usher syndrome type 2D (Ebermann *et al*., 2007; Mathur *et al*., 2015). The existing knowledge of *WHRN* transcript isoforms has predominantly been derived from research utilizing mutant mouse models. These studies have identified four *WHRN* transcript isoforms expressed in the mouse inner ear: two full-length isoforms featuring different exon compositions, an N-terminal, and a C-terminal isoform (Belyantseva *et al*., 2005; Ebrahim *et al*., 2016; Mburu *et al*., 2003). In contrast, the current understanding of *WHRN* isoforms expressed in the human retina is limited. While previous studies by Mburu *et al*. (2003) and Belyantseva *et al*. (2005) have suggested the presence of human retinal *WHRN* transcript isoforms based on cDNA clones, the study by van Wijk *et al*. (2006) verified the presence of two *WHRN* transcript isoforms in the human retina using semi-quantitative RT-PCR analysis: the full-length isoform (ENST00000362057.4) and a C-terminal isoform (ENST0000674048.1) (Figure 4A).

**Figure 4:**
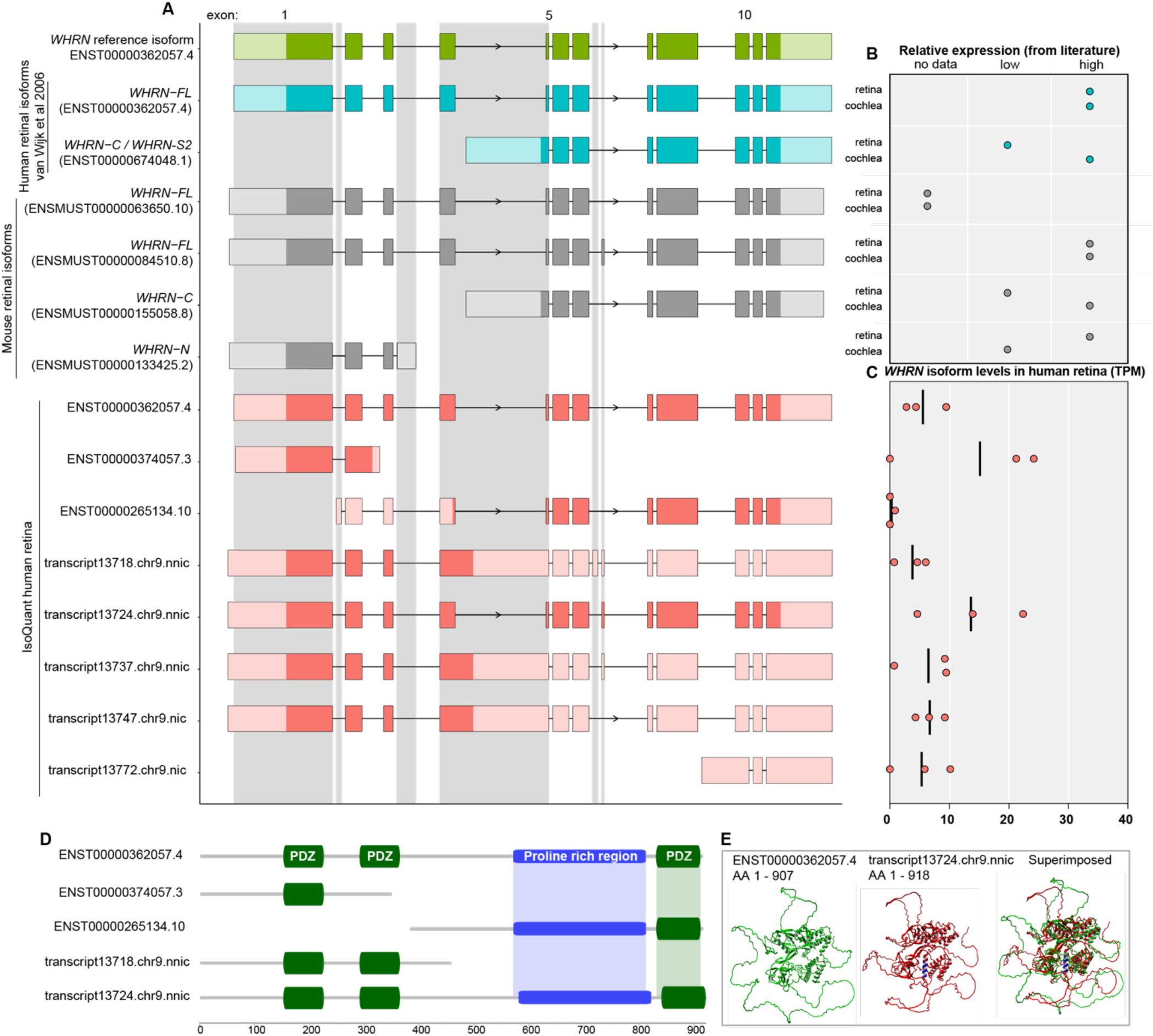
*WHRN* transcript isoforms identified by IsoQuant analysis compared to known isoforms from the literature. **A.** The GENCODE reference transcript is depicted at the top in green, followed by human *WHRN* transcript isoforms from literature in blue (van Wijk *et al*., 2006) and the murine transcript isoforms in grey (Belyantseva *et al*., 2005; Ebrahim *et al*., 2016; Mburu *et al*., 2003). The *WHRN* IsoQuant transcripts are depicted in red. The light green, blue, grey and red colors indicate the untranslated regions (UTR) and the dark green, blue, grey and red colors indicate the open reading frame (ORF) of each transcript. Differences between the IsoQuant transcripts and the GENCODE reference transcript are highlighted in grey boxes. **B.** Relative expression of *WHRN* isoforms based on literature in either the regna or the cochlea. **C.** The Transcripts Per Million (based on dataset 1) for each IsoQuant transcript isoform are presented for the three individual samples. **D.** The predicted 2D protein domain architecture of the encoded WHRN protein isoforms. Light blue and green boxes highlight the difference between the WHRN reference isoform and the protein isoform encoded by exon 7B-containg transcript13724.chr9.nic. **E.** AlphaFold2 3D protein predictions of two WHRN isoforms; reference isoform ENST00000362057.4 in green and transcript13724.chr9.nic in red, with the alpha helix encoded by the novel exon 7B highlighted in blue.

The IsoQuant analysis confirmed the expression of the full-length reference isoform of the *WHRN* gene in the human retina; however, none of the predicted transcript isoforms corresponded to the previously reported C-terminal isoform, ENST0000674048.1. While this finding contradicts the semi-quantitative RT-PCR analysis by van Wijk *et al*. (2006), it aligns with studies that show the absence of a C-terminal isoform in the murine retina, as it is reported to be expressed exclusively in the mouse inner ear (Mathur and Yang, 2015). Notably, the IsoQuant analysis identified an alternative C-terminal *WHRN* transcript isoform starting with a distinct non-coding exon (ENST00000265134.10), although its expression is relatively low (0.3 ± 0.5 TPM) (Figure 4C).

Based on studies in the mouse retina, the presence of an N-terminal *WHRN* isoform was suggested for the human retina as well (Belyantseva *et al*., 2005; Mburu *et al*., 2003). The human isoforms ENST0000374057.3 (*WHRN-N*) and ENST0000699486 (*WHRN-NT2*) are thought to resemble the mouse N-terminal *WHRN* transcript isoforms, but have never been experimentally validated (and thus not shown as reference in Figure 4A). However, IsoQuant results indicated the presence of the N-terminal transcript isoform ENST00000374057.3. While this transcript is only detected in 2 out of the 3 retinal samples, the observed TPM levels (15.1 ± 13.2) in those two samples indicate this isoform is expressed at high levels.

Additionally, multiple IsoQuant transcript isoforms suggested the presence of transcripts with intron 4 retention. This intron retention introduces a premature stop codon, and therefore these transcripts are predicted to encode a truncated protein containing only the first two PDZ domains (Figure 4D). Manual curation of sequenced reads also confirmed intron 4 retention in ∼50% of the sequenced reads. Similarly, intron 4 retention was observed in ONT long-read mRNA sequencing data obtained from three independent retinal samples. While such transcripts would typically be disregarded due to the early termination of translation, the fact that nearly half of the predicted transcripts exhibit intron 4 retention indicates this alternative transcript isoform may warrant further investigation for potential functional relevance.

Finally, two novel exons were identified within intron 7: designated exon 7A and 7B. While exon 7A was exclusively observed as a non-coding exon and always co-occurred with exon 7B, exon 7B was additionally detected as an independent protein-coding exon in transcript13724.chr9.nic. Manual inspection of the sequencing data also revealed the widespread presence of exon 7B in the Iso-Seq transcripts, which was further corroborated by an independent ONT long-read mRNA sequencing. This 33-nucleotide-long exon 7B is not present in the GENCODE reference annotations for the human *WHRN* gene but does resemble a corresponding 33-nucleotide exon found in the mouse *Whrn* consensus sequence. Consequently, transcript13724.chr9.nic matches the full-length mouse *Whrn* transcript, which contains 13 protein-coding exons, in contrast to the 12-exon human *WHRN* reference sequence. Structural modeling with AlphaFold2 revealed that this novel exon 7B encodes an additional alpha helix in of 11 amino acids in the center of the protein (Figure 4E).

Comparing the TPM levels of the *WHRN* reference transcript (5.5 ± 3.5 TPM for ENST00000362057.4) with those of the exon 7B-containing transcript (13.6 ± 8.9 TPM for transcript13724.chr9.nnic), suggests that the exon 7B-containing transcript could be the predominant retinal isoform. Because biallelic pathogenic variants in exon 7B could potentially cause USH2D, we queried WGS data of our in-house cohort of unsolved IRD patients (de Bruijn *et al*., 2023), but found no candidate pathogenic variants in this exon.

### Identifying variable *USH2A* transcription start sites, pseudoexon-prone regions and the successful enrichment of full-length isoform B transcripts using Samplix Xdrop Sort

The *USH2A* gene is one of the most frequently mutated genes among those associated with IRDs. It exhibits a diverse mutational profile, with a higher incidence of missense variants observed in cases of non-syndromic autosomal recessive retinitis pigmentosa (arRP), while truncating variants are more commonly associated with the syndromic cases of Usher syndrome type 2A (Lenassi *et al*., 2014). The *USH2A* gene gives rise to two distinct protein isoforms. Isoform A is the shorter variant, composed of 1546 amino acids encoded by a 21-exon mRNA transcript (Eudy *et al*., 1998). Conversely, isoform B represents the larger isoform, composed of 5202 amino acids and encoded by a transcript consisting of 72 exons (Van Wijk *et al*., 2004).

The PacBio standard- and long transcript workflows were unable to capture the full-length *USH2A* transcripts encoding isoform B. Similarly, we did not identify isoform B transcripts in an independent ONT long-read mRNA sequencing dataset. However, IsoQuant analysis confirmed the presence of shorter isoform A transcripts. Additionally, our IsoQuant analysis revealed transcripts (transcript51429.chr1.nnic and transcript51430.chr1.nnic) that demonstrate significant diversity in their 5’ transcription start sites, leading to the inclusion of additional amino acids at the 5’ end of the encoded open reading frame (ORF), while maintaining the same reading frame and sharing the same 3’ UTR as isoform A (Figure 5A). Notably, this diversity in transcription start sites was also observed in the independent ONT long-read mRNA sequencing data.

**Figure 5:**
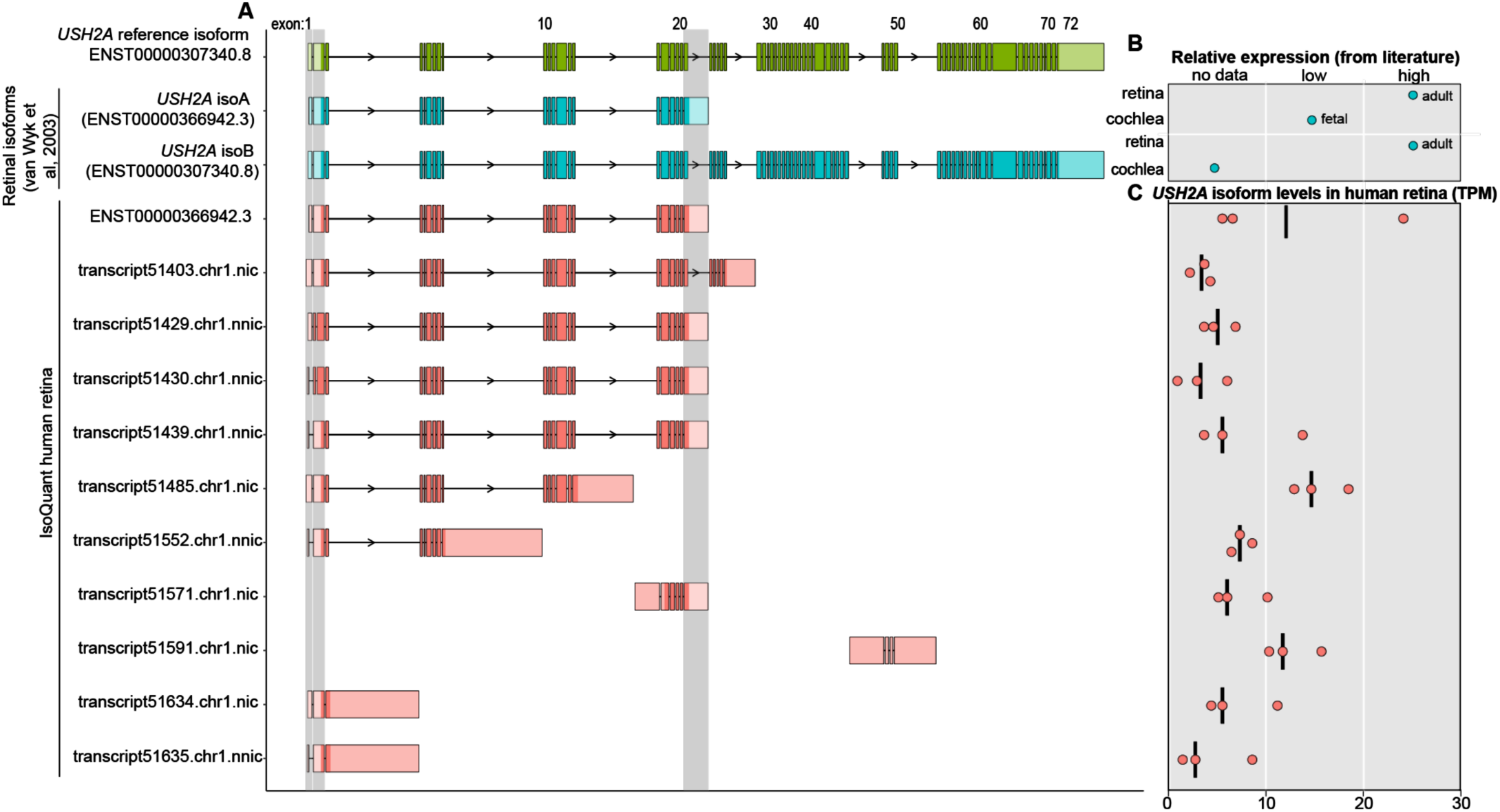
*USH2A* transcript isoforms identified by IsoQuant analysis compared to known transcripts from the literature. **A.** The GENCODE reference transcript is depicted at the top in green, followed by the known human *USH2A* transcript isoforms in blue (Van Wijk *et al*., 2004). The *USH2A* IsoQuant transcripts are depicted in red. The light green, blue and red colors indicate the untranslated regions (UTR) and the dark green, blue and red colors indicate the open reading frame (ORF) of each transcript. Differences between the IsoQuant transcript isoforms and the GENCODE reference transcript are highlighted in grey boxes. **B.** Relative expression of *USH2A* isoforms based on literature in either the regna or the cochlea. **C.** The Transcripts Per Million (based on dataset 1) for each IsoQuant transcript are presented for the three individual samples.

IsoQuant analysis revealed that the most abundant transcript is spanning exons 1 through 15 (transcript51485.chr1.nic), as evidenced by the TPM values (15.35 ± 2.85 TPM; Figure 5C). Manual inspection of the sequenced reads using the BAM files in IGV confirmed substantial read support for this alternative isoform. However, the 3’ end of this transcript is followed by a stretch of 12 adenine nucleotides, which may have led to internal priming of the oligo(dT) primer. Moreover, the shorter length of this transcript isoform compared to transcripts that encode known usherin isoforms may have led to its preferential amplification and loading on the PacBio SMRT-cells, potentially explaining its higher TPM values. Therefore, it remains to be determined whether this transcript corresponds to a *bona fide* transcript isoform or an experimental artifact.

Several N-terminal *USH2A* transcripts were identified that spanned different exon ranges, including exons 1-9 (transcript 51552.chr1.nnic), 1-26 (transcript51403.chr1.nic), and 1-3 (transcript51634.chr1.nic), as well as a transcript covering exons 16-21 (transcript 51591.chr1.nic) (Figure 5). Given the limitations of the PacBio sequencing workflows in fully sequencing *USH2A* transcripts encoding usherin isoform B, and the observation of numerous partial reads upon manual inspection of the BAM files, these factors may have contributed to the IsoQuant transcript models of shorter, partial *USH2A* transcript isoforms. Hence, the authenticity of these transcript models remains uncertain, as they could represent genuine transcripts or merely partial reads.

The value of the optimized long transcript workflow is clearly demonstrated through the manual examination of the sequenced reads (Figure 6). While tiling of reads resulting from the PacBio standard workflow samples did not fully cover the complete *USH2A* isoform B transcript, the reads from the sample prepared following the optimized long transcript workflow together were able to cover all 72 exons of *USH2A.* These data did not reveal any evidence of natural exon skipping (NES); a phenomenon where specific exons are excluded from the mature transcripts in healthy tissues, as observed for the *ABCA4* gene by Tomkiewicz (2024). Furthermore, numerous reads across all samples support the presence of isoform A transcripts and the observed variation in 5’ transcription start sites. The individual reads do not clearly indicate whether this 5’ variation is specific to isoform A or is also a feature of isoform B. However, we identified reads extending beyond exon 21 that are linked to the variation in 5’ start site, indicating that this 5’ variation may also be present in the larger isoform B transcripts.

**Figure 6:**
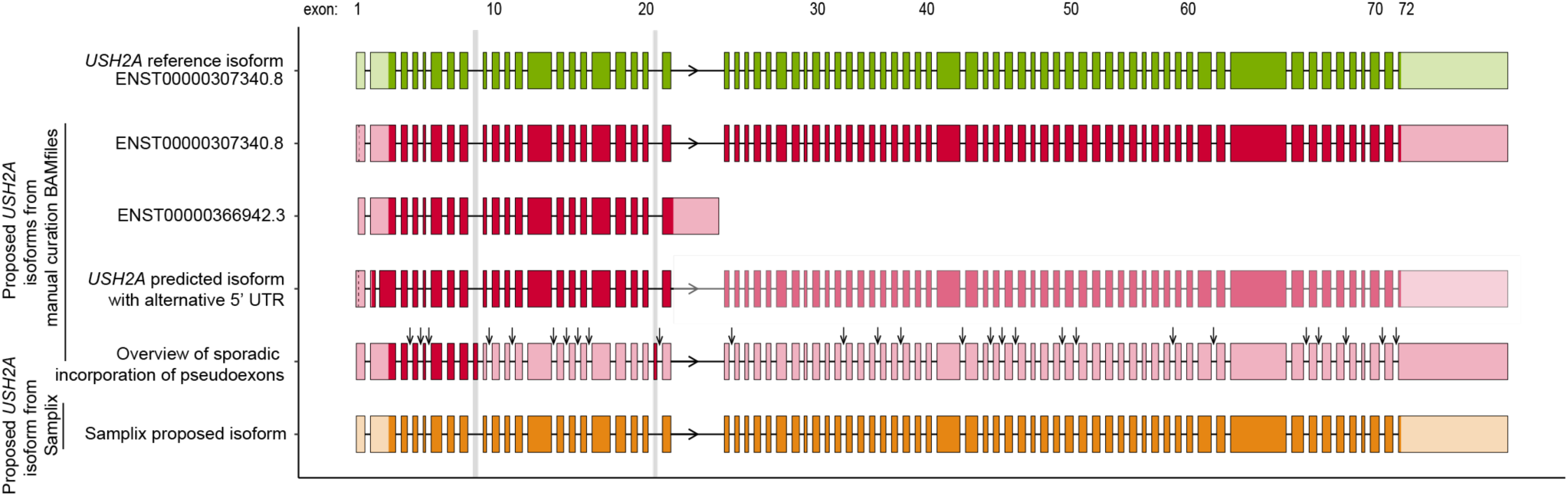
*USH2A* proposed transcript isoforms from manual curation and Samplix Xdrop targeted enrichment. The GENCODE reference transcript is depicted at the top in green, followed by the proposed *USH2A* transcript isoforms and events based on manual curation of BAM-files using the Integrative Genomics Viewer (IGV) in red, and proposed transcript isoforms following the Samplix Xdrop targeted enrichment in orange. The light green, red and orange colors indicate the untranslated regions (UTR) and the dark green, red and orange colors indicate the open reading frame (ORF) of each transcript. Differences between the proposed transcript isoforms and the GENCODE reference transcript are highlighted in grey boxes. The overview of sporadic incorporation of cryptic exons indicates the presence of PE8 and PE20 as previously described by Reurink *et al*. (2023). Additionally, locatons where cryptic exons are occasionally incorporated at sites that are not yet noted in literature are highlighted by the overlaying black arrows in the overview.

Manual curation of sequenced transcripts further uncovered sporadic inclusion of intronic sequences, for example the inclusion of 87 bp of intron 20, previously identified as pseudoexon 20 (PE20) by Reurink *et al*. (2023). In addition to the previously reported *USH2A* PE8 and PE20 identified by Reurink *et al*. (2023) our investigation also revealed the occasional incorporation of other previously uncharacterized cryptic exons. The intronic regions where these cryptic exons are observed are denoted by arrows in the overview provided in Figure 6, and genomic positions are provided in table S7. As demonstrated by Reurink *et al*. (2023), pathogenic deep-intronic variants can result in the inclusion of PEs harboring an in-frame stop codon across all *USH2A* transcripts. This prompted us to evaluate WGS data from our in-house cohort of unsolved IRD patients (de Bruijn *et al*., 2023). However, we did not identify any candidate pathogenic deep-intronic variants surrounding the identified sporadic cryptic exons. Although we failed to detect additional pathogenic variants in our cohort that could result in the inclusion of these incidental cryptic exons across all *USH2A* transcripts, the observed sporadic inclusion of cryptic exons corresponding to PE8 and PE20 suggests that the other identified cryptic exons may highlight regions susceptible to pathogenic PE-inducing variants.

To overcome the limitations of the PacBio standard and long-transcript workflows, as well as the ONT long-read mRNA sequencing of independent retina samples, in sequencing the full-length *USH2A* isoform B transcripts, we employed a targeted enrichment approach on cDNA using the Samplix Xdrop Sort technology in an effort to capture these *USH2A* transcripts. The Samplix Xdrop Sort is a technique designed for single-molecule enrichment of genomic DNA. Excitingly, we demonstrated the utility of this approach to capture cDNA as well. By using three detection sequences targeting either 5’ - middle and 3’ sites of the longest known transcript encoding usherin isoform B, we were able to enrich for *USH2A* transcripts, as evidenced by the qPCR results (Table S6). The enriched pool of transcripts was then subjected to HiFi long-read sequencing and subsequent cDNA analysis. Manual curation of the obtained sequencing reads revealed that the detection sequence targeting exons 30-31 enabled us to capture and sequence complete, full-length transcripts encoding usherin isoform B for the first time. This provides conclusive evidence for the presence of full-length *USH2A* transcripts encoding isoform B in the human retina (Figure 6).

### Iso-Seq reveals a novel coding *ADGRV1* exon, but lacks the capability to sequence full-length transcripts encoding the VLGR1b isoform

With a coding sequence spanning 19.6 kb, *ADGRV1* - previously known as *MASS1, VLGR1*, and *GPR98* - is the largest Usher-associated gene, and variants in this gene are responsible for Usher syndrome type 2C (Weston *et al*., 2004). To date, three distinct *ADGRV1* transcript isoforms have been identified in human: *VLGR1a*, *VLGR1b*, and *VLGR1c*. *VLGR1b* represents the transcript encoding the longest isoform, composed of 90 exons and encoding a protein of 6,306 amino acids. The C-terminal *VLGR1a* and N-terminal *VLGR1c* transcript isoforms arise from alternative splicing events. Specifically, *VLGR1a* is generated by a 5’ extension of exon 65, introducing a novel open reading frame and producing a 1,976 amino acid protein. *VLGR1c* results from an alternative splice donor site within exon 31 resulting in an 83 bp truncation at the 3’ end of exon 31, which shifts the reading frame and introduces a premature stop codon. The relative expression of these *ADGRV1* transcript isoforms in the retina and cochlea remains unknown, as evidence for these transcripts has primarily been derived from brain tissue studies (McMillan *et al*., 2002). Similarly, the known murine *Adgrv1* transcript isoforms have also been predominantly characterized in brain rather than cochlear or retinal tissues (Skradski *et al*., 2001; Yagi *et al*., 2005).

Similar challenges to those encountered with the large coding sequence of *USH2A* were observed for *ADGRV1*, which prevented the PacBio standard- and long transcript workflows and ONT long-read mRNA sequencing, from capturing the transcript encoding the longest isoform VLGR1b. Surprisingly, the IsoQuant analysis did not identify any transcript isoforms corresponding to the shorter *VLGR1a* and *VLGR1c* transcripts, despite their smaller sizes not posing a barrier to sequencing (Figure 7A). However, manual curation of the sequenced transcripts revealed that approximately half of the reads contained the 83 bp truncation at the 3’ end of exon 31, as also observed in the N-terminal *VLGR1c* transcript isoform. While no sequencing reads spanned exon 1 to exon 31, the observed utilization of the splice site resulting in the 83 bp truncation of exon 31 suggests the presence of *VLGR1c* transcripts in the human retina (Figure 8). Additionally, the 5’ extension of exon 65, which defines the start of the *VLGR1a* transcript, was observed in reads across all sequenced samples. Notably, only the sample prepared following the PacBio long transcript workflow contained reads that perfectly spanned from the 5’ extension of exon 65 to exon 90, demonstrating the presence of the *VLGR1a* transcript in the human retina. Further manual inspection of the PacBio Iso-Seq data and independent ONT long-read mRNA sequencing data revealed that nearly half of the reads contained an in-frame novel exon situated in intron 39, which has been designated as exon 39A (33 bp in size). However, the precise relationship between this novel exon and the specific *ADGRV1* transcript isoforms remains unclear. Because biallelic pathogenic variants in exon 39A could potentially cause USH2C, we queried WGS data of our in-house cohort of unsolved IRD patients (de Bruijn *et al*., 2023), but found no candidate pathogenic variants in this exon. Collectively, these findings underscore the value of thorough manual curation of sequencing data, combined with the long transcript workflow enrichment approach, for identifying novel events and *ADGRV1* transcripts in the human retina.

**Figure 7:**
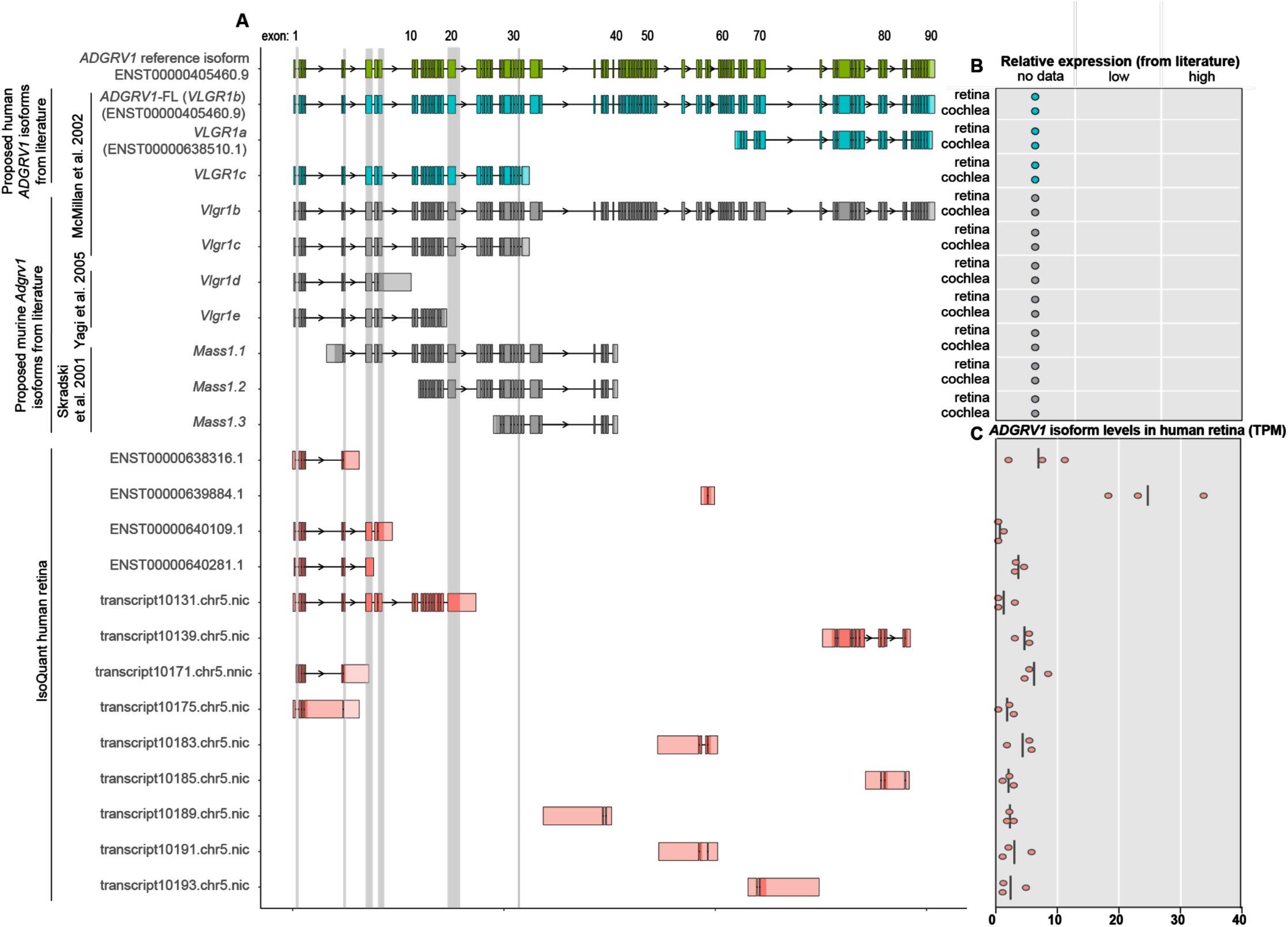
*ADGRV1* transcript isoforms identified by IsoQuant analysis compared to known isoforms from the literature. **A.** The GENCODE reference transcript is depicted at the top in green, followed by the known human *ADGRV1* isoforms in blue (McMillan *et al*., 2002), and the murine isoforms in grey (McMillan *et al*., 2002; Skradski *et al*., 2001; Yagi *et al*., 2005). The *ADGRV1* IsoQuant transcripts are depicted in red. The light green, blue, grey and red colors indicate the untranslated regions (UTR) and the dark green, blue, grey and red colors indicate the open reading frame (ORF) of each transcript. Differences between the IsoQuant transcript isoforms and the GENCODE reference transcript are highlighted in grey boxes. **B.** Relative expression of *ADGRV1* isoforms based on literature in either the regna or the cochlea. **C.** The Transcripts Per Million (based on dataset 1) for each IsoQuant isoform are presented for the three individual samples.

While the observations for the *VLGR1a* and *VLGR1c* transcripts necessitated manual curation of the sequenced transcripts in IGV, and were not part of the IsoQuant output, the IsoQuant analysis did identify the presence of isoforms that are analogous to those previously reported in mice. Specifically, the identified transcript ENST00000640109.1, spanning exons 1-9 and terminating in intron 9, closely resembles the murine *Vlgr1d* transcript (Yagi *et al*., 2005). Similarly, the IsoQuant transcript10131.chr5.nic, spanning exons 1-20, exhibits a high degree of similarity to the murine *Vlgr1e* transcript (Yagi *et al*., 2005), with the exception that the murine isoform ends in intron 19. While both predicted transcript isoforms demonstrate relatively low TPM values, they were also identified through manual curation of sequenced transcripts. In contrast, neither the IsoQuant analysis nor the manual curation provided any indication of the presence of isoforms similar to the murine *Mass 1.1, Mass 1.2*, and *Mass 1.3* transcript isoforms.

The IsoQuant analysis identified ENST00000639884.1 as the most abundant transcript, but this isoform only encompasses exons 57 and 58. Interestingly, manual curation of transcripts revealed no individual reads supporting the presence of this short transcript, raising questions about the reason for its prediction. Furthermore, IsoQuant identified multiple transcripts terminating in intron 6, such as ENST00000638316.1, which were corroborated by manual curation across all sequenced samples. Similarly, the IsoQuant transcript10139.chr5.nic was also supported by manual curation. In contrast, while the IsoQuant algorithm identified several other short transcripts with very small open reading frames (transcript101.83.chr5.nic, transcript101.85.chr5.nic, transcript101.89.chr5.nic, transcript101.91.chr5.nic and transcript101.93.chr5.nic), identical reads were not identified through the manual curation of sequenced reads, questioning the biological relevance of these transcripts. The limitations of PacBio Iso-Seq in sequencing *ADGRV1* transcripts encoding the full-length VLGR1b isoform, coupled with the manual curation findings of numerous partial reads, may have contributed to the IsoQuant-based identification of shorter, incomplete *ADGRV1* transcript isoforms. Consequently, the authenticity of these predicted isoforms remains uncertain, as they could represent genuine transcripts or simply reflect incomplete sequence data.

In an effort to capture full-length *ADGRV1* (19.6 kb) transcripts encoding the longest isoform VLGR1b, we employed the targeted enrichment approach using the Samplix Xdrop Sort system, by utilizing three detecting sequences targeting the 5’, middle, and 3’ regions of *VLGR1b* transcripts. qPCR results confirmed the enrichment of *ADGRV1* transcripts (Table S6). While we cannot conclude that the detection sequences were able to capture the full-length isoform, manual curation of sequenced reads revealed that the 5’ detection sequence (targeting exons 16-17) provided the best coverage, spanning exons 3 to 73 and 79 to 90. This suggests the existence of the full-length *ADGRV1* isoform encoding VLGR1b in the human retina. Although the absence of reads mapping against exons 74 – 78 in the 5’ enriched sample may hint at natural exon skipping (NES), no reads were found that indicated that exon 73 is immediately followed by exon 79. Furthermore, we did not observe any indication for missing exons 74 -78 in the data obtained with the PacBio standard- and long transcript workflows. Therefore, we assume that the lack of coverage for exons 74-78 is due to the Samplix Xdrop amplification procedure of multiple displacement amplification (MDA), which introduces branched transcripts. Finally, 5’ target analysis confirmed transcripts similar to the murine *Vlgr1e* isoform in the human retinal transcriptome (Figure 8).

**Figure 8:**
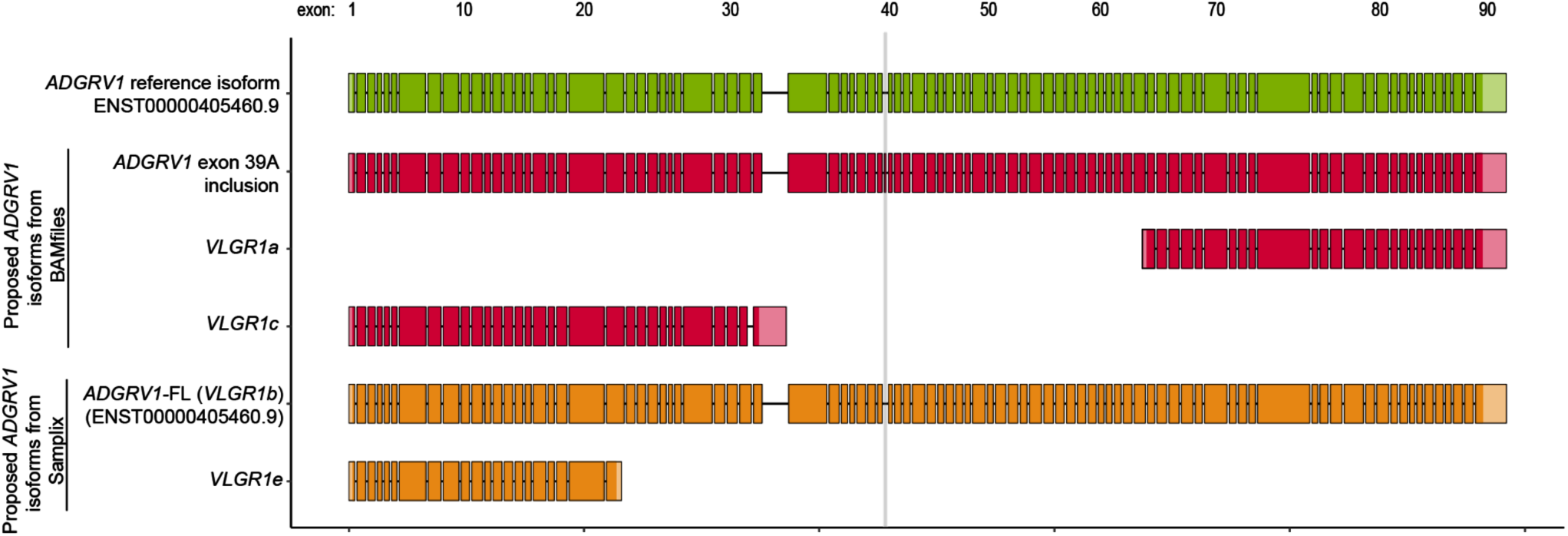
*ADGRV1* proposed transcript isoforms from manual curation and Samplix Xdrop targeted enrichment. The GENCODE reference transcript is depicted at the top in green, followed by the *ADGRV1* proposed transcript isoforms and events based on manual curation of BAM-files using the Integrative Genomics Viewer (IGV) in red, and proposed transcript isoforms following the Samplix Xdrop targeted enrichment in orange. The light green, red and orange colors indicate the untranslated regions (UTR) and the dark green, red and orange colors indicate the open reading frame (ORF) of each transcript. Differences between the proposed transcript isoforms and the GENCODE reference transcript are highlighted in grey box.

## DISCUSSION

Given the limitations of current sequencing approaches in comprehensively characterizing the largest transcripts expressed in the human neural retina, we conducted PacBio long-read RNA sequencing following the standard library preparation and an optimized workflow to enrich for long transcripts in the human neural retina. Our goal was to characterize the transcript isoforms associated with Usher syndrome, as the 11 related genes are known to produce large transcripts up to 19.6 kb in size. While the range of CCS reads obtained in this study is consistent with other published PacBio Iso-Seq analyses (Leung *et al*., 2021), and our optimized workflow enabled sequencing of transcripts up to 15 kb, this was insufficient for Usher syndrome-associated genes *USH2A* and *ADGRV1*, with transcripts of 18.9 kb and 19.6 kb, respectively. With a targeted enrichment using the Samplix Xdrop System we successfully captured full-length 18.9 kb *USH2A* transcripts. By integrating algorithmic analysis and manual curation of sequenced transcripts, we have identified several novel transcript isoforms that may exhibit unique functionalities in the human retina. Additionally, the discovery of previously unidentified exons could provide diagnostic value. Although querying whole-genome sequencing data from our internal cohort of unsolved IRD cases (de Bruijn *et al*., 2023) did not uncover any candidate pathogenic variants within these newly discovered exons, these genomic regions could still be incorporated when screening for disease-causing variants associated with Usher syndrome. Furthermore, the identification of regions with incidental inclusion of cryptic exons may highlight potential hotspots for deep-intronic pathogenic variants associated with Usher syndrome (Reurink *et al*., 2023). Collectively, these results provide valuable insights into retinal Usher syndrome-associated transcript isoforms, with implications for improving diagnostics and therapeutic development.

The detailed analysis of the 11 Usher syndrome-associated genes was conducted using our previously published PacBio long-read mRNA Iso-Seq dataset, which was generated following the PacBio standard workflow (Riepe *et al*., 2024). This analysis was further supplemented with a PacBio Iso-Seq dataset of an additional sample prepared using an optimized long transcript workflow, aimed at capturing longer transcripts. A comparison of the PacBio standard workflow and our optimized long transcript workflow demonstrated that the optimized workflow, which generated transcripts up to 15 kb with an average size of 4.7 kb, significantly surpasses the standard workflow’s average read length of 2.6 kb. This enhancement is particularly advantageous for capturing the longer coding sequences characteristic for several Usher syndrome-associated genes. However, incorporating the long transcript workflow dataset into a combined IsoQuant analysis with the standard workflow datasets presented challenges in accurately classifying transcript isoforms using the IsoQuant algorithm. Including the long transcript dataset in the combined analysis resulted in fewer FSMs, a shift in the number of NIC, and more NNIC transcripts being predicted from the standard workflow datasets compared to analyzing them independently. This suggests IsoQuant’s sensitivity to input data characteristics, particularly read length and intron retention, which were more prevalent in the long transcript dataset. Manual curation of sequenced reads confirmed a higher occurrence of intron-retaining transcripts in the long transcript dataset. While this workflow enriched for larger transcripts, it also appears to have captured more transcripts where splicing is ongoing, resulting in transcripts with retained introns. This could distort the IsoQuant algorithm’s annotation of isoforms, complicating the identification of fully spliced transcript isoforms. Although we previously (Riepe *et al*., 2024) and in our current analysis preferred using IsoQuant over other analysis tools like SQANTI3 and TALON due to its lower rate of false-positive isoforms (Pardo-Palacios *et al*., 2024; Prjibelski *et al*., 2023), we anticipated that IsoQuant would provide more precise isoform predictions when analyzing the standard- and long-transcript datasets separately rather than in a combined analysis. This separation allows for more accurate transcript isoform predictions and comparisons by maintaining the consistency of input data characteristics within each analysis set. In conclusion, the optimized long transcript workflow significantly improves the detection of large transcripts, but it necessitates careful consideration of its impact on isoform classification algorithms. Future studies should refine these algorithms or develop approaches to leverage both standard and long transcript workflows effectively.

Our focus on the identification of Usher syndrome-associated transcripts in the human retina enabled a more detailed examination compared to genome-wide screens and discoveries that rely heavily on advanced tools and algorithms to manage large-scale data. By zooming in on Usher genes, we were able to sift through novel transcripts with greater precision through manual analysis. Therefore, our integrated approach of combining algorithmic analysis with the manual curation of sequenced transcripts enabled the identification of novel isoforms and alternative splicing events across the 11 Usher syndrome-associated genes, which we highlighted for *MYO7A, WHRN, USH2A*, and *ADGRV1*. For instance, we discovered a novel predominant *MYO7A* transcript isoform. Previous research using a targeted PCR approach had only identified two retinal *MYO7A* transcript isoforms differing in exon 35 length, with the truncated exon 35 variant being the predominant form (Gilmore *et al*., 2023). This PCR-based targeted methodology is incapable of identifying isoforms characterized by potentially novel transcription initiation and termination sites. In contrast, the PacBio Iso-Seq approach allowed us to identify a novel *MYO7A* transcript characterized by an alternative 5’ start site paired with an extended 3’ untranslated region. While Gilmore *et al*. (2023) have suggested that both previously known *MYO7A* transcript isoforms should be considered when designing gene therapies, our findings indicate that the isoforms with novel transcription 5’ start site were the most predominant in the human retina. Therefore, it might be even more important to incorporate the novel *MYO7A* transcript isoforms we have identified in the design of retinal gene augmentation therapies. Furthermore, the novel, previously unannotated *MYO7A* transcription start site could be incorporated into diagnostic screening pipelines as it may harbor pathogenic variants of potential clinical significance. Additionally, we observed that the utilization of this alternative 5’ start site for the *MYO7A* transcript isoform is associated with an extended 3’ untranslated region. Previous studies have indicated that the selection of alternative polyadenylation (APA) sites is coupled with alternative splicing patterns (Anvar *et al*., 2018; Zhang *et al*., 2019). The length of the 3’-UTR can influence mRNA stability, the accessibility of regulatory elements like miRNA binding sites, and the efficiency of protein translation. Furthermore, recent findings suggest that the choice of a 3’ end site is broadly influenced by the site of transcription initiation (Alfonso-Gonzalez *et al*., 2023). This implies that the utilization of this alternative 5’ start site for the *MYO7A* transcript may have significant regulatory implications for gene expression and protein function within the human neural retina.

Additionally, sporadic intron retention events were observed across all 11 Usher syndrome-associated genes, likely indicating ongoing splicing in these transcripts. Notably, the high frequency of intron 30 and 37 retention in the *MYO7A* transcripts is particularly interesting. It is well-established that the order of intron removal during splicing does not follow a consecutive 5’-3’ direction, and the speed of intron removal is not dependent on intron length (Pandya-Jones and Black, 2009; Singh and Padgett, 2009). However, Gazzoli *et al*. (2016) provided evidence for multi-step splicing of dystrophin (*DMD*) introns, demonstrating that slowly spliced introns may lead to the formation of “exon blocks” - groups of consecutive exons that tend to be spliced together as a unit. This can occur when the flanking introns are inefficiently spliced, resulting in the exons being treated as a single unit during intermediate steps of the final mature RNA. Similarly, the observed retention of introns 30 and 37 in the *MYO7A* transcripts may be indicative of the generation of exon blocks for exons 31-37.

The interplay between intron retention and exon blocks may particularly be relevant for the design of antisense oligonucleotide (ASO) therapies, for example when aiming to induce the exclusion of exons harboring pathogenic variants. Gazzoli *et al*. (2016) speculate that skipping multiple exons, such as exons 45-51 in the *DMD* gene, would be beneficial for 40% of Duchenne patients. However, as the number of exons requiring exclusion increases, the simultaneous removal of all target exons from a single pre-mRNA molecule becomes more difficult. This difficulty arises because multiple ASOs must concurrently bind to their respective exons (Echigoya *et al*., 2018). To address this challenge, Gazzoli *et al*. (2016) hypothesize that targeting the entire *DMD* exon block 45-51, rather than individual exons, could be achieved with fewer ASO molecules. With several ASO-based splicing modulation therapies under development for Usher syndrome-associated RP, some also based on multiple exon skipping (Schellens *et al*., 2023), investigating the potential presence of exon blocks is a worthwhile endeavor. Our current dataset could help uncover additional exon blocks within other Usher genes.

Another notable instance of intron retention was observed in *WHRN* transcripts, where both the computational IsoQuant analysis and manual examination of the sequencing data revealed that nearly half of the sequenced *WHRN* transcripts exhibited retention of intron 4. While intron retention was previously thought to disrupt protein production due to the introduction of premature stop codons that trigger nonsense-mediated decay (NMD), research by Boutz *et al*. (2015) has identified a class of intron-retaining transcripts that are polyadenylated, sequestered in the nucleus, and resistant to NMD. These retained introns can undergo subsequent splicing and export to the cytoplasm for translation, potentially enabling cells to rapidly adapt to environmental changes or stress. Although the RNA used to generate the PacBio Iso-Seq libraries was extracted from total human neural retina samples, and we cannot conclusively determine if the *WHRN* transcripts containing intron 4 retention originate from the nucleus, the widespread occurrence of this *WHRN* intron 4 retention variant across the analyzed samples suggests it may hold functional significance and could potentially be governed by a similar regulatory mechanism as the retained introns described by Boutz *et al*. (2015).

The PacBio Iso-Seq standard and long-transcript workflows were unable to sequence transcripts encoding the currently known largest isoforms of usherin and ADGRV1, but did identify the presence of *USH2A* transcripts encoding usherin isoform A, and several shorter *ADGRV1* transcripts analogous to murine *Vlgr1d* and *Vlgr1e*. Moreover, the long-transcript workflow provided improved coverage across these genes. Manual examination of the *USH2A* reads revealed the sporadic inclusion of intronic sequences, which we have termed cryptic exons. The occurrence of these sporadic cryptic exons may be a result of ongoing splicing, as previous studies using ultra-deep sequencing have demonstrated that recursive splicing can take place in which introns are spliced out in multiple steps (Gazzoli *et al*., 2016). In this process, a canonical donor 5’ splice site may be spliced to an internal acceptor site, generating a 5’ ratcheting point from the juxtaposed exon and the 5’ splice site sequences, and when a similar mechanism occurs at the 3’ splice site, this can lead to the generation of a cryptic exon as an intermediate product (Gazzoli *et al*., 2016). Alternatively, the identification of cryptic exons may hint at regions susceptible to pathogenic pseudoexon-inducing variants. For instance, manual inspection of the *USH2A* reads revealed the sporadic inclusion of 87 bp of intron 20, previously identified as PE20 by Reurink *et al*. (2023). Their study also reported sporadic inclusion of PE20 in *USH2A* transcripts. However, they found that the c.4397-3890A>G variant resulted in consistent inclusion of PE20, containing an in-frame stop codon (p.Ala1465_Ala1466ins*5), across all *USH2A* transcripts. An ASO-based splicing correction therapy was developed to target this aberrant PE inclusion. This example illustrates how deep intronic variants can lead to the permanent inclusion of potentially pathogenic PEs. The observed sporadic inclusion of a cryptic exon corresponding to PE20 suggests that other cryptic exons we identified might indicate regions susceptible to pathogenic PE-inducing variants. This underscores the diagnostic potential of a detailed Iso-Seq data analysis, which could inform the development of targeted diagnostic panels and therapeutic strategies.

Despite using our PacBio long-transcript workflow, we were unable to sequence the currently known largest transcript isoforms of *USH2A* and *ADGRV1*. We chose to utilize PacBio Iso-Seq for our study due to its reputation for offering higher sequencing accuracy compared to Oxford Nanopore Technologies (ONT) (Tvedte *et al*., 2021). While the ONT platform presents advantages, such as adaptive sampling for target enrichment and direct RNA or cDNA sequencing, which may improve transcript capture in future studies, an independent ONT dataset from human retina samples also failed to capture the full-length transcripts for *USH2A* and *ADGRV1*, highlighting the current challenges faced by both sequencing platforms. Considering these challenges, we utilized the Samplix Xdrop Sort system for an ‘indirect target enrichment’ (Madsen *et al*., 2020) on human retina cDNA to enrich for *USH2A* and *ADGRV1* transcripts. This facilitated the successful capture and sequencing of *ADGRV1* transcripts as well as cDNA molecules of *USH2A* encoding the largest isoform B, thereby demonstrating the presence of this complete cDNA molecule (18.9 kb) for the first time. In addition to its original purpose of genomic DNA enrichment, this marks the first demonstration of applying the Samplix Xdrop Sort system to enrich cDNA samples. However, a limitation of this technique is that the captured cDNA molecules require an amplification step using multiple displacement amplification (MDA), which introduces branched transcripts. Consequently, an enzymatic digestion step is necessary prior to sequencing. Ideally, the workflow would enable full-length amplification of the captured molecules followed by PacBio long-read sequencing, but this remains an area for future improvement.

## CONCLUSION AND FUTURE DIRECTIONS

Our findings underscore the importance of employing integrated sequencing and analysis approaches to capture the full complexity of the transcriptome in specialized tissues like the human neural retina. Through a comprehensive analysis of Usher syndrome-associated transcript isoforms in the human retina, we revealed a more intricate isoform landscape than previously understood. This analysis led to the discovery of novel isoforms and splicing events, with significant implications for diagnostics and therapeutic development. This work exemplifies the benefits of combining algorithmic and detailed manual analyses to achieve a deeper understanding. We advocate for the utilization of the human neural retina sequencing datasets we generated here for similar comprehensive studies, as these datasets have the potential to yield valuable insights into other genes and significantly advance our understanding of complex transcriptomic landscapes.

Future research should focus on further refining sequencing and enrichment techniques to enhance the capture of full-length transcripts for large genes. Additionally, functional studies are necessary to validate the transcript isoforms and elucidate their roles in retinal physiology and pathology. This comprehensive understanding will be instrumental in developing effective genetic therapies to prevent vision loss in individuals with Usher syndrome.

## DATA AVAILABILITY

The PacBio Iso-Seq data generated for this study, and the ONT long-read mRNA sequencing data used for validation of selected transcripts and events, are publicly accessible through the European Genome-Phenome Archive at www.ega-archive.org, under the accession number EGAD50000000720. Additionally, genome browser tracks of the analyzed PacBio Iso-Seq data can be accessed at euro.ucsc.edu/s/tabeariepe/retina_atlas. The original codes have been made publicly accessible through the GitHub repository (https://github.com/erikdevrieze/USH_retina).

## ACKNOWLEDGEMENTS

This study was financially supported by Stichting UitZicht (2019-16), CUREUsher and Stichting Ushersyndroom. Purchase of the Samplix Xdrop Sort Instrument was enabled by an internal Radboudumc technology innovation grant (to AH). We would like to thank the R&D department of Samplix ApS for their expert guidance in optimizing the research protocol to enable cDNA capturing. Finally, we also thank the Radboud Technology Center Genomics for the library preparation and sequencing of all samples.

## REFERENCES

Alfonso-Gonzalez C, Legnini I, Holec S, Arrigoni L, Ozbulut HC, Mateos F, Koppstein D, Rybak-Wolf A, Bonisch U, Rajewsky N et al. 2023. Sites of transcription initiation drive mRNA isoform selection. Cell 186(11):2438–2455 e22.

Anvar SY, Allard G, Tseng E, Sheynkman GM, de Klerk E, Vermaat M, Yin RH, Johansson HE, Ariyurek Y, den Dunnen JT et al. 2018. Full-length mRNA sequencing uncovers a widespread coupling between transcription initiation and mRNA processing. Genome Biol 19(1):46.

Belyantseva IA, Boger ET, Naz S, Frolenkov GI, Sellers JR, Ahmed ZM, Griffith AJ, Friedman TB. 2005. Myosin-XVa is required for tip localization of whirlin and differential elongation of hair-cell stereocilia. Nat Cell Biol 7(2):148–56.

Boutz PL, Bhutkar A, Sharp PA. 2015. Detained introns are a novel, widespread class of post-transcriptionally spliced introns. Genes Dev 29(1):63–80.

Braun TA, Mullins RF, Wagner AH, Andorf JL, Johnston RM, Bakall BB, Deluca AP, Fishman GA, Lam BL, Weleber RG et al. 2013. Non-exomic and synonymous variants in ABCA4 are an important cause of Stargardt disease. Hum Mol Genet 22(25):5136–45.

Cao H, Wu J, Lam S, Duan R, Newnham C, Molday RS, Graziotto JJ, Pierce EA, Hu J. 2011. Temporal and tissue specific regulation of RP-associated splicing factor genes PRPF3, PRPF31 and PRPC8--implications in the pathogenesis of RP. PLoS One 6(1):e15860.

Ciampi L, Mantica F, Lopez-Blanch L, Permanyer J, Rodriguez-Marin C, Zang J, Cianferoni D, Jimenez-Delgado S, Bonnal S, Miravet-Verde S et al. 2022. Specialization of the photoreceptor transcriptome by Srrm3-dependent microexons is required for outer segment maintenance and vision. Proc Natl Acad Sci U S A 119(29):e2117090119.

de Bruijn SE, Rodenburg K, Corominas J, Ben-Yosef T, Reurink J, Kremer H, Whelan L, Plomp AS, Berger W, Farrar GJ et al. 2023. Optical genome mapping and revisiting short-read genome sequencing data reveal previously overlooked structural variants disrupting retinal disease-associated genes. Genet Med 25(3):100345.

Ebermann I, Scholl HP, Charbel Issa P, Becirovic E, Lamprecht J, Jurklies B, Millan JM, Aller E, Mitter D, Bolz H. 2007. A novel gene for Usher syndrome type 2: mutations in the long isoform of whirlin are associated with retinitis pigmentosa and sensorineural hearing loss. Hum Genet 121(2):203–11.

Ebrahim S, Ingham NJ, Lewis MA, Rogers MJC, Cui R, Kachar B, Pass JC, Steel KP. 2016. Alternative Splice Forms Influence Functions of Whirlin in Mechanosensory Hair Cell Stereocilia. Cell Rep 15(5):935–943.

Echigoya Y, Lim KRQ, Nakamura A, Yokota T. 2018. Multiple Exon Skipping in the Duchenne Muscular Dystrophy Hot Spots: Prospects and Challenges. J Pers Med 8(4).

Eudy JD, Weston MD, Yao S, Hoover DM, Rehm HL, Ma-Edmonds M, Yan D, Ahmad I, Cheng JJ, Ayuso C et al. 1998. Mutation of a gene encoding a protein with extracellular matrix motifs in Usher syndrome type IIa. Science 280(5370):1753–7.

Gazzoli I, Pulyakhina I, Verwey NE, Ariyurek Y, Laros JF, t Hoen PA, Aartsma-Rus A. 2016. Non-sequential and multi-step splicing of the dystrophin transcript. RNA Biol 13(3):290–305.

Gilmore WB, Hultgren NW, Chadha A, Barocio SB, Zhang J, Kutsyr O, Flores-Bellver M, Canto-Soler MV, Williams DS. 2023. Expression of two major isoforms of MYO7A in the retina: Considerations for gene therapy of Usher syndrome type 1B. Vision Res 212:108311.

Guilford P, Ayadi H, Blanchard S, Chaib H, Le Paslier D, Weissenbach J, Drira M, Petit C. 1994. A human gene responsible for neurosensory, non-syndromic recessive deafness is a candidate homologue of the mouse sh-1 gene. Hum Mol Genet 3(6):989–93.

Gustavsson EK, Zhang D, Reynolds RH, Garcia-Ruiz S, Ryten M. 2022. ggtranscript: an R package for the visualization and interpretation of transcript isoforms using ggplot2. Bioinformatics 38(15):3844–3846.

Hasson T, Heintzelman MB, Santos-Sacchi J, Corey DP, Mooseker MS. 1995. Expression in cochlea and retina of myosin VIIa, the gene product defective in Usher syndrome type 1B. Proc Natl Acad Sci U S A 92(21):9815–9.

Jumper J, Evans R, Pritzel A, Green T, Figurnov M, Ronneberger O, Tunyasuvunakool K, Bates R, Zidek A, Potapenko A et al. 2021. Highly accurate protein structure prediction with AlphaFold. Nature 596(7873):583–589.

Konagurthu AS, Whisstock JC, Stuckey PJ, Lesk AM. 2006. MUSTANG: a multiple structural alignment algorithm. Proteins 64(3):559–74.

Krieger E, Vriend G. 2014. YASARA View - molecular graphics for all devices - from smartphones to workstations. Bioinformatics 30(20):2981–2.

Lenassi E, Saihan Z, Bitner-Glindzicz M, Webster AR. 2014. The effect of the common c.2299delG mutation in USH2A on RNA splicing. Exp Eye Res 122:9–12.

Letunic I, Khedkar S, Bork P. 2021. SMART: recent updates, new developments and status in 2020. Nucleic Acids Res 49(D1):D458–D460.

Leung SK, Jeffries AR, Castanho I, Jordan BT, Moore K, Davies JP, Dempster EL, Bray NJ, O’Neill P, Tseng E et al. 2021. Full-length transcript sequencing of human and mouse cerebral cortex identifies widespread isoform diversity and alternative splicing. Cell Rep 37(7):110022.

Li H. 2021. New strategies to improve minimap2 alignment accuracy. Bioinformatics 37(23):4572–4574.

Li S, Mecca A, Kim J, Caprara GA, Wagner EL, Du TT, Petrov L, Xu W, Cui R, Rebustini IT et al. 2020. Myosin-VIIa is expressed in multiple isoforms and essential for tensioning the hair cell mechanotransduction complex. Nat Commun 11(1):2066.

Liu MM, Zack DJ. 2013. Alternative splicing and retinal degeneration. Clin Genet 84(2):142–9.

Liu X, Vansant G, Udovichenko IP, Wolfrum U, Williams DS. 1997. Myosin VIIa, the product of the Usher 1B syndrome gene, is concentrated in the connecting cilia of photoreceptor cells. Cell Motil Cytoskeleton 37(3):240–52.

Madsen EB, Hoijer I, Kvist T, Ameur A, Mikkelsen MJ. 2020. Xdrop: Targeted sequencing of long DNA molecules from low input samples using droplet sorting. Hum Mutat 41(9):1671–1679.

Mathur P, Yang J. 2015. Usher syndrome: Hearing loss, retinal degeneration and associated abnormalities. Biochim Biophys Acta 1852(3):406–20.

Mathur PD, Zou J, Zheng T, Almishaal A, Wang Y, Chen Q, Wang L, Vashist D, Brown S, Park A et al. 2015. Distinct expression and function of whirlin isoforms in the inner ear and retina: an insight into pathogenesis of USH2D and DFNB31. Hum Mol Genet 24(21):6213–28.

Mburu P, Mustapha M, Varela A, Weil D, El-Amraoui A, Holme RH, Rump A, Hardisty RE, Blanchard S, Coimbra RS et al. 2003. Defects in whirlin, a PDZ domain molecule involved in stereocilia elongation, cause deafness in the whirler mouse and families with DFNB31. Nat Genet 34(4):421–8.

McMillan DR, Kayes-Wandover KM, Richardson JA, White PC. 2002. Very large G protein-coupled receptor-1, the largest known cell surface protein, is highly expressed in the developing central nervous system. J Biol Chem 277(1):785–92.

Murphy D, Cieply B, Carstens R, Ramamurthy V, Stoilov P. 2016. The Musashi 1 Controls the Splicing of Photoreceptor-Specific Exons in the Vertebrate Retina. PLoS Genet 12(8):e1006256.

Niyadurupola N, Sidaway P, Osborne A, Broadway DC, Sanderson J. 2011. The development of human organotypic retinal cultures (HORCs) to study retinal neurodegeneration. Br J Ophthalmol 95(5):720–6.

Osborne A, Hopes M, Wright P, Broadway DC, Sanderson J. 2016. Human organotypic retinal cultures (HORCs) as a chronic experimental model for investigation of retinal ganglion cell degeneration. Exp Eye Res 143:28–38.

Pandya-Jones A, Black DL. 2009. Co-transcriptional splicing of constitutive and alternative exons. RNA 15(10):1896–908.

Pardo-Palacios FJ, Wang D, Reese F, Diekhans M, Carbonell-Sala S, Williams B, Loveland JE, De Maria M, Adams MS, Balderrama-Gutierrez G et al. 2024. Systematic assessment of long-read RNA-seq methods for transcript identification and quantification. Nat Methods 21(7):1349–1363.

Prjibelski AD, Mikheenko A, Joglekar A, Smetanin A, Jarroux J, Lapidus AL, Tilgner HU. 2023. Accurate isoform discovery with IsoQuant using long reads. Nat Biotechnol 41(7):915–918.

Reurink J, Weisschuh N, Garanto A, Dockery A, van den Born LI, Fajardy I, Haer-Wigman L, Kohl S, Wissinger B, Farrar GJ et al. 2023. Whole genome sequencing for USH2A-associated disease reveals several pathogenic deep-intronic variants that are amenable to splice correction. HGG Adv 4(2):100181.

Riepe T, Stemerdink M, Salz R, Duenas Rey A, de Bruijn SE, Boonen E, Tomkiewicz TZ, Kwint M, Gloerich J, Wessels HJ. 2024. A proteogenomic atlas of the human neural retina. bioRxiv:2024.05. 22.595273.

Ruiz-Ceja KA, Capasso D, Pinelli M, Del Prete E, Carrella D, di Bernardo D, Banfi S. 2023. Definition of the transcriptional units of inherited retinal disease genes by meta-analysis of human retinal transcriptome data. BMC Genomics 24(1):206.

Sarantopoulou D, Brooks TG, Nayak S, Mrcela A, Lahens NF, Grant GR. 2021. Comparative evaluation of full-length isoform quantification from RNA-Seq. BMC Bioinformatics 22(1):266.

Schellens RTW, Broekman S, Peters T, Graave P, Malinar L, Venselaar H, Kremer H, De Vrieze E, Van Wijk E. 2023. A protein domain-oriented approach to expand the opportunities of therapeutic exon skipping for USH2A-associated retinitis pigmentosa. Mol Ther Nucleic Acids 32:980–994.

Singh J, Padgett RA. 2009. Rates of in situ transcription and splicing in large human genes. Nat Struct Mol Biol 16(11):1128–33.

Skradski SL, Clark AM, Jiang H, White HS, Fu YH, Ptacek LJ. 2001. A novel gene causing a mendelian audiogenic mouse epilepsy. Neuron 31(4):537–44.

Tomkiewicz TZ. 2024. Skipping, elongation, and restoration. A tale of ABCA4 splicing to pave the road towards therapeutic applications. Radboud University Nijmegen.

Tvedte ES, Gasser M, Sparklin BC, Michalski J, Hjelmen CE, Johnston JS, Zhao X, Bromley R, Tallon LJ, Sadzewicz L et al. 2021. Comparison of long-read sequencing technologies in interrogating bacteria and fly genomes. G3 (Bethesda) 11(6).

Udovichenko IP, Gibbs D, Williams DS. 2002. Actin-based motor properties of native myosin VIIa. J Cell Sci 115(Pt 2):445–50.

Van Wijk E, Pennings RJ, te Brinke H, Claassen A, Yntema HG, Hoefsloot LH, Cremers FP, Cremers CW, Kremer H. 2004. Identification of 51 novel exons of the Usher syndrome type 2A (USH2A) gene that encode multiple conserved functional domains and that are mutated in patients with Usher syndrome type II. The American Journal of Human Genetics 74(4):738–744.

van Wijk E, van der Zwaag B, Peters T, Zimmermann U, Te Brinke H, Kersten FF, Marker T, Aller E, Hoefsloot LH, Cremers CW et al. 2006. The DFNB31 gene product whirlin connects to the Usher protein network in the cochlea and retina by direct association with USH2A and VLGR1. Hum Mol Genet 15(5):751–65.

Wang B, Tseng E, Regulski M, Clark TA, Hon T, Jiao Y, Lu Z, Olson A, Stein JC, Ware D. 2016. Unveiling the complexity of the maize transcriptome by single-molecule long-read sequencing. Nat Commun 7:11708.

Weil D, Blanchard S, Kaplan J, Guilford P, Gibson F, Walsh J, Mburu P, Varela A, Levilliers J, Weston MD et al. 1995. Defective myosin VIIA gene responsible for Usher syndrome type 1B. Nature 374(6517):60–1.

Weston MD, Luijendijk MW, Humphrey KD, Moller C, Kimberling WJ. 2004. Mutations in the VLGR1 gene implicate G-protein signaling in the pathogenesis of Usher syndrome type II. Am J Hum Genet 74(2):357–66.

Yagi H, Takamura Y, Yoneda T, Konno D, Akagi Y, Yoshida K, Sato M. 2005. Vlgr1 knockout mice show audiogenic seizure susceptibility. J Neurochem 92(1):191–202.

Zhang Z, So K, Peterson R, Bauer M, Ng H, Zhang Y, Kim JH, Kidd T, Miura P. 2019. Elav-Mediated Exon Skipping and Alternative Polyadenylation of the Dscam1 Gene Are Required for Axon Outgrowth. Cell Rep 27(13):3808–3817 e7.

