## supplemental data for "Pushing the limits of single molecule transcript sequencing to uncover the largest disease-associated transcript isoforms in the human neural retina"

**SUPPLEMENTS:**

**Table S1: Details of human neural retina samples used for PacBio Iso-Seq**

| Sample ID | Sex | Age | Time until enucleation | RIN value |
|---|---|---|---|---|
| Sample 1 | Male | 59 | 2 hours 35 minutes | 8.2 |
| Sample 2 | Male | 63 | 9 hours 5 minutes | 7.3 |
| Sample 3 | Female | 58 | 11 hours 55 minutes | 7.5 |

**Table S2: Composition of *USH2A* and *ADGRV1* detection sequence and validation sequence primers.**

| Primer ID | Forward sequence | Reverse Sequence | Targeted exons |
|---|---|---|---|
| *USH2A* Front dPCR | ACAACTGAGACTGCTGTTAACC | GACCATGGCACTGACATCTC | 8-9 |
| *USH2A* Front qPCR | GCAAAGCAAACGTTATTGGGCT | TACACTGGCAGGGCTCACAT | 12-13 |
| *USH2A* Middle dPCR | GAAAGCCTATAGTGAGGACAGCA | TCTCAGTACAGCCAGCCAAA | 30-31 |
| *USH2A* Middle qPCR | TGTTTCGGTTGGTTGCCTCC | GGTCCAGGTTTGTCCTCTGC | 40-41 |
| *USH2A* End dPCR | TGGAGTGACACCTTCCTCCT | CCTTCGTCAGTCGTGCAGAT | 68-69 |
| *USH2A* End qPCR | ACTGTGGGAAGCCATCATGG | TGTCTGTGAATGTGGTGCGT | 71-72 |
| *ADGRV1* Front dPCR | ACTGAATGGCACTGGAGGAG | AGTTGGCAGTCTCACCTTCC | 13-14 |
| *ADGRV1* Front qPCR | TGCTAGCAAGATTGGATGGGA | ACAATGGTGGCACTTCCCAA | 16-17 |
| *ADGRV1* Middle dPCR | GGCTTTGTCTTCAGCCAATG | GCTGGTGTGAAGGAGTGGAT | 50-51 |
| *ADGRV1* Middle qPCR | CTCCTTCACACCAGCCTCAG | CGCTCCGAATTCCAGCAGTA | 49-50 |
| *ADGRV1* End dPCR | ATAAAGTGGACGTGGTGCCA | AAACGTCCACCGACAAGCAT | 70-71 |
| *ADGRV1* End qPCR | GCAACATGACCCCAACACTG | CAACCAGGGCTAGGAAACGT | 72-73 |

**Table S3: Details of human neural retina samples used for Oxford Nanopore Technology sequencing.**

| Sample ID | Time until enucleation | RIN value | Reads (passed) |
|---|---|---|---|
| Sample 1 | < 20 hours | > 8 | 10,206,706 |
| Sample 2 | < 20 hours | > 8 | 18,463,865 |
| Sample 3 | < 20 hours | > 8 | 11,388,116 |

**Table S4: *MYO7A* qPCR primer sequences**

| Primer ID | Forward sequence | Reverse Sequence | Targeted exons |
|---|---|---|---|
| *MYO7A* 5' canonical start | CTTCCTGAGTCCTCCGTGC | CACATGGTCCCCCTGCTG | 1-2 |
| *MYO7A* 5' alternative start | ACCAGGCGTTCAAGACCTAC | CATCATCCACCACCTGGACC | Alt.1-2 |
| *MYO7A* 3' end | ACGTCACTGGGCTACAAGATG | CTCCCCGTGACTGTTCACTT | 48-49 |
| *GUSB* | AGAGTGGTGCTGAGGATTGG | CCCTCATGCTCTAGCGTGTC | 2-3 |

### TABLE S5: OVERVIEW OF OBSERVED EVENTS FOLLOWING MANUAL CURATION OF READS IN BAM FILES SAMPLES 1-4.

| **Gene** | **Curated transcripts*** | **Novel exons** (in-frame)** | **Novel exon** (out of frame)** | **In-frame skipping events** | **Out of frame skipping** events | **Alternative/novel splice junctions** | **Intron retentions** |
|---|---|---|---|---|---|---|---|
| *MYO7A* | ENST00000409709.9 | Alternative transcription start site 5' of canonical exon 3, Exon 30A, 31A, 31B | Exon 2A, 2B, 2C, 2D, 4A, 15A, 27A, 44A, 46B | Exon 3, 11, 26 | Exon 9, (9+10 combined), 14 | 5' extension of exon 8, 12, 17, 28, 42, 44<br><br>3' extension of exon 12<br><br>5' truncation of exon 35, 40, 44, 16 | Intron 8, 9, 11, 28, 30 (**observed in ~25% of reads**), 33, 37 (**observed in ~25% of reads**), 38, 40, 41, 42, 43, 44 |
| *USH1C* | ENST00000005226.12 | Exon 15A, 20A | Exon 15B, 18A, alternative penultimate exon | Exon 2, 11, 15 | Exon 19, 20 | 5' extension of exons 6, 12, 16<br><br>3' extension of exon 5<br><br>3' truncation of exon 14 | Intron 3, 4, 7, 10, (10 + 11 combined), (22+23 combined), 23, 23 (with internal fragment spliced out), 24 (with internal fragment spliced out) |
| *CDH23* | ENST00000224721.12 | Exon 11A, 31A, 66A | Exon 1A, 25A, 26A, 44A, 44B, 44C, 45A, 48A, 48B, 48C, 48D | Exon 12, 34, 46, (56-57 combined skip), **exon 69 skip observed in ~80% of reads** | - | 5' extension of exons 6, 12, 46, 49, 55, 60, 65, 70<br><br>3' extension of exons 16, 55, 60, 63<br><br>5' truncation of exons 58, 60<br><br>3' truncation of exons 58, 59, 60 | Intron 7, 23 (45-46 combined), 46, 50, 64, 65, (65-66 combined) |
| *PCDH15* | ENST0000064397 | Exon 1A, 2A, 5A, 9A, 9B, 18A, 18B | Exon 1B, 9C, 9D, 9E, 13A, 13B, 15A, 16A, 17A, 18C, 26A, 26B, 26C, 32A, 32B, 36A | Exon 19, 21, 22 27, 31, 33 (exon 33, 34, 35, 36 combined skip) | Exon 10, 14, 16, 17 23, 26 | 5' truncation of exon 11<br><br>3' truncation of exon 36 | - |
| *SANS* | ENST00000614341.5 | - | - | - | - | - | - |
| *CIB2* | ENST00000258930.8 | - | - | - | - | - | - |
| *USH2A* | ENST00000307340.8 | Exon 3A | Exons 4A, 5A, 8A, 9A, 9B, 9C, 11A, 11B, 11C, 12A, | Exon 8, (exon 6,7,8 combined skip) 28, | Exon 3, (exon 5, 6, 7, 8, 9 combined skip) exon 6, 9, 12, (exon | 5' extension of exon 65, 70<br><br>3' extension of exon 14, 15 | Intron 5, 6 |

| | | | | | | | |
|---|---|---|---|---|---|---|---|
| | | | 13A, 13B, 13C, 14A, 15A, 16A, 20A, 20B, 20C, 20D, 22A, 32A, 32B, 35A, 37A, 41A, 43A, 44A, 44B, 45A, 49A, 50A, 58A, 61A, 64A, 64B, 65A, 67A, 70A, 71A | (exon 33-37 combined skip) | 12-13 combined skip), exon 14, (exon 15-19 combined skip), (exon 15-22 combined skip) exon 26, (exon 33-34 combined skip) , exon 42, 52, 55, (exon 65-67 combined skip), | 5’ truncation of exon 8, 19, 46, 64<br><br>3’ truncation of exon 1, 6, 10, 13, 25, 46, | |
| *ADGRV1* | ENST00000405460.9 | Exon 1A, **39A inclusion observed in ~50% of reads**, 86A | Exon 77A, 77B, 77C, 79A, 79B, 86A | Exon 4 | Exon 67, 76, (exon 76-77 combined skip), (exon 88-89 combined skip) | 5’ extension of exon 28, 58, 89<br><br>3’ extension of exon 2, 25, 43, 58, 69, 88<br><br>5’ truncation of exon 26, 42, 52<br><br>3’ truncation of exon 6, 49 | Intron 4, 39, 52, 56, 64, 88 |
| *WHRN* | ENST00000362057.4 | Exon 2A, **7B inclusion observed in ~80% of reads** (7B + 7G combined) | (Exon 7A + 7B combined), 7C, 7D 7E, 7F, 7G, 7H | Exon 3, 7 | Exon 4, (exon 5+6 combined skip) | 5’ extension of exon 5<br><br>3’ extension of exon 4<br><br>5’ truncation of exon 7, 10<br><br>3’ truncation of exon 9 | **Intron 4 retention observed in ~50% of reads**<br><br>Intron 6, (combined intron 4 + 9 retention) |
| *CLRN1* | ENST00000327047.6 | - | - | - | - | - | - |
| *ARSG* | ENST00000621439.5 | Exon 8A, 10A, alternative terminal exon | Alternative transcription start site 5’ of canonical exon 1, Exon 2A, 10B, 10C, 10D | - | (Exon 7+8 combined skip), (exon 9+10 combined skip) | 5’ extension exon 9<br><br>3’ extension exon 2, 3, 5, 6, 11, | - |

*Curated MANE select transcripts that were used to annotate observed events.

** Novel identified exons are designated with alphabetical labels (e.g., 'A', 'B') to denote their relative positioning. For instance, '30A' indicates a novel exon observed downstream of the canonical exon 30.

**Table S6: Mean qPCR CT values of cDNA samples before and after Samplix Xdrop Sort**

| Target | Mean CT value unenriched sample | Mean CT value enriched sample |
|---|---|---|
| *USH2A* 5' Front | 25.8 | 13.4 |
| *USH2A* Middle | 25.7 | 15.7 |
| *USH2A* 3' End | 23.7 | 17.9 |
| *ADGRV1 5'* Front | 26.4 | 16.3 |
| *ADGRV1* Middle | 24.5 | 16.5 |
| *ADGRV1* 3' End | 26.1 | 12.2 |

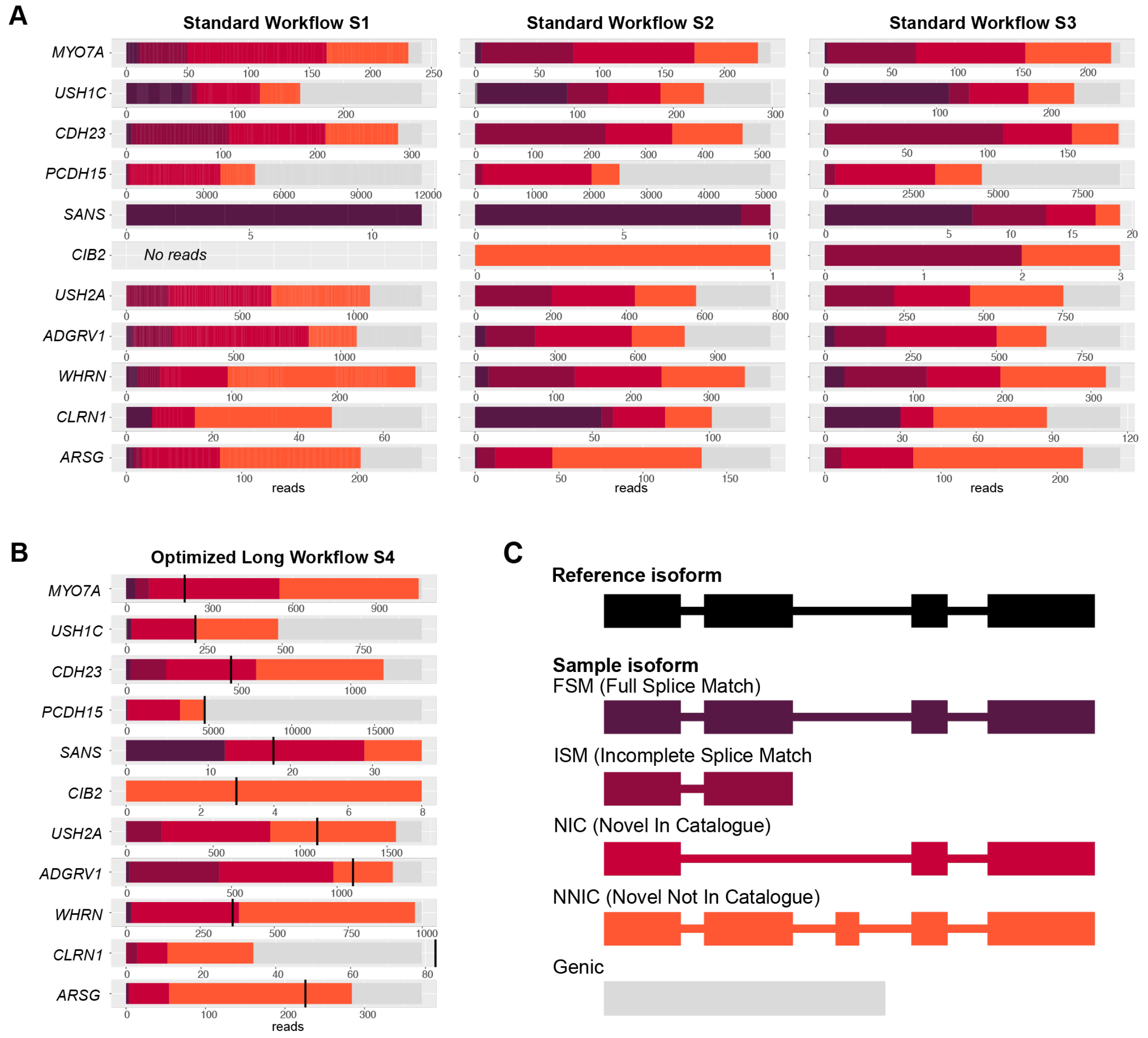


**Figure S1. Comparison between Usher syndrome isoform transcripts obtained from PacBio standard workflow and optimized long workflow datasets. A:** Number of reads for each USH gene distributed per isoform category: Full Splice Match (FSM), Incomplete Splice Match (ISM), Novel In Catalogue (NIC) and Novel Not In Catalogue (NNIC). S1, S2 and S3 represent the individual retina donor samples, for which libraries were prepared according to the PacBio standard workflow. **B:** Number of reads for each USH gene distributed per isoform category. S4 represents the sample prepared following an optimized PacBio long workflow. **C:** Schematic representation that compares retinal transcripts to the GENCODE reference transcriptome: Full Splice Matches (FSMs), Incomplete Splice Matches (ISM), Novel In Catalog (NIC) and Novel Not In Catalog (NNIC). Genomic contamination is indicated in grey.

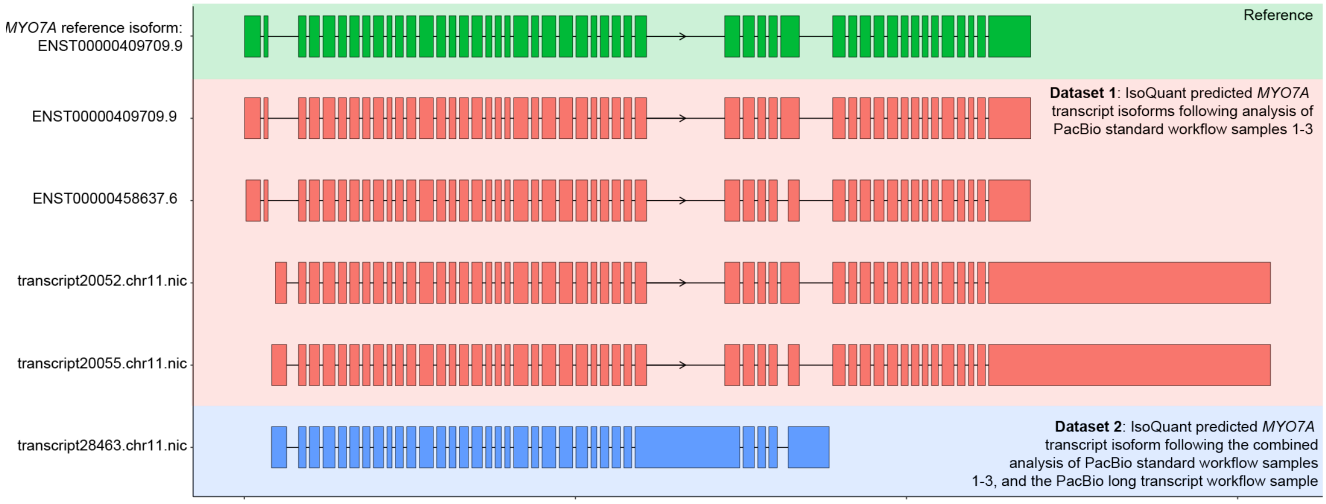


**Figure S2. Examination of IsoQuant algorithm outputs across different datasets, highlighting the influence of including or excluding the PacBio long transcript workflow sample on the IsoQuant-predicted *MYO7A* isoforms.** The *MYO7A* gene serves as an example to illustrate the divergence in identified isoforms by the IsoQuant algorithm. The green annotation represents the *MYO7A* reference isoform. In red are the *MYO7A* isoforms resulting from the IsoQuant analysis of standard workflow samples 1-3, as identified in dataset 1 (Riepe et al., 2024). This includes both Full Splice Matches (FSMs) to the reference isoform, as well as Novel In Catalogue (NICs) and Novel Not In Catalogue (NNICs) isoforms. In blue is the *MYO7A* isoform following the IsoQuant analysis of the combined standard and long transcript workflow samples in dataset 2. Despite the increased input data, this analysis only yielded a single NNIC isoform, with no FSMs or NICs detected.

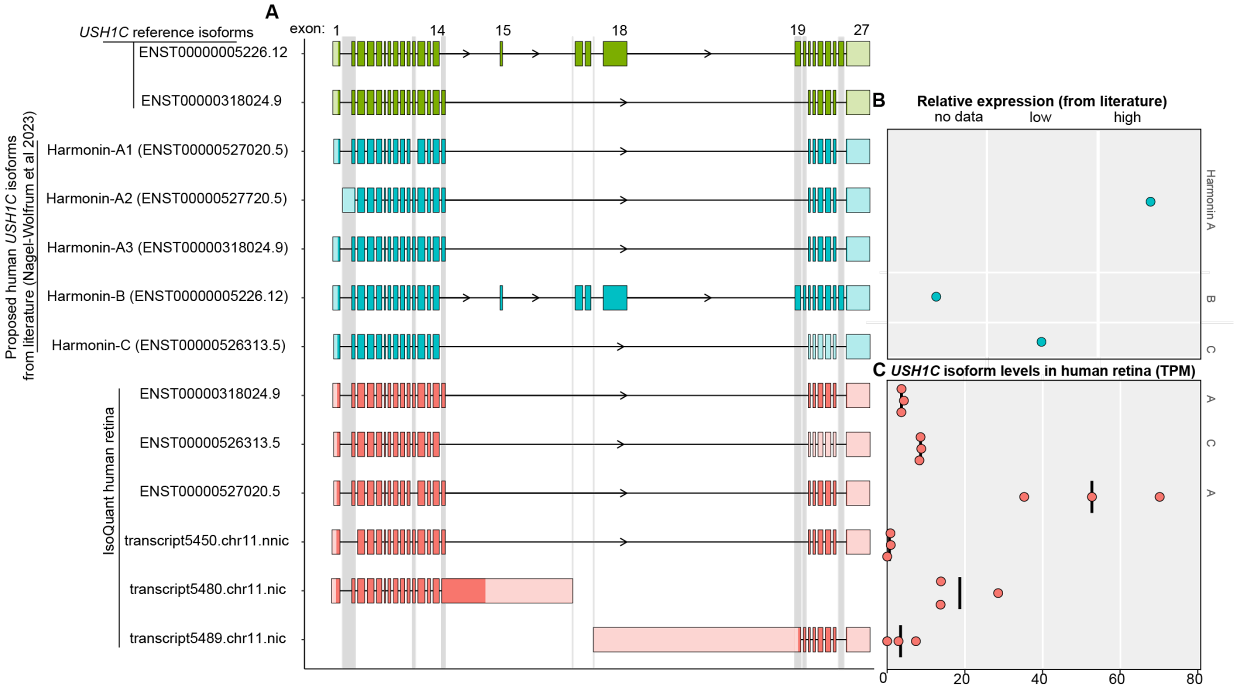


**Figure S3: *USH1C* isoforms identified by IsoQuant analysis compared to known isoforms from the literature. A.** The GENCODE reference transcripts are depicted at the top in green, followed by the known human *USH1C* isoforms in blue (Nagel-Wolfrum et al., 2023). The *USH1C* IsoQuant transcripts are depicted in red. The light green, blue and red colors indicate the untranslated regions (UTR) and the dark green, blue and red colors indicate the open reading frame (ORF) of each transcript. Differences between the IsoQuant isoforms and the GENCODE reference transcript are highlighted in grey boxes. IsoQuant predicts the absence of the Harmonin-B isoform in the human retina. **B.** Relative expression of *USH1C* isoforms based on literature in either the retina or the cochlea. **C.** The Transcripts Per Million (based on dataset 1) for each IsoQuant identified isoform are presented for the three individual samples.

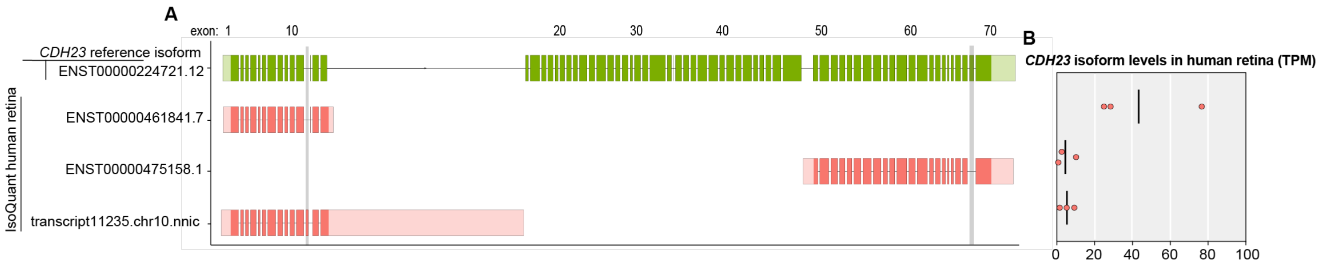


**Figure S4: *CDH23* isoforms identified by IsoQuant analysis compared to known isoform. A.** The GENCODE reference transcript is depicted at the top in green, followed by *CDH23* IsoQuant transcripts depicted in red. The light green and red colors indicate the untranslated regions (UTR) and the dark green and red colors indicate the open reading frame (ORF) of each transcript. Differences between the IsoQuant isoforms and the GENCODE reference transcript are highlighted in grey boxes. IsoQuant analysis indicates skipping of exon 69, which is in line with findings of Siemens et al., (2002) that indicate the presence of this penultimate exon in inner ear transcripts, and its absence in other tissues such as the retina. **B** The Transcripts Per Million (based on dataset 1) for each IsoQuant identified isoform are presented for the three individual samples.

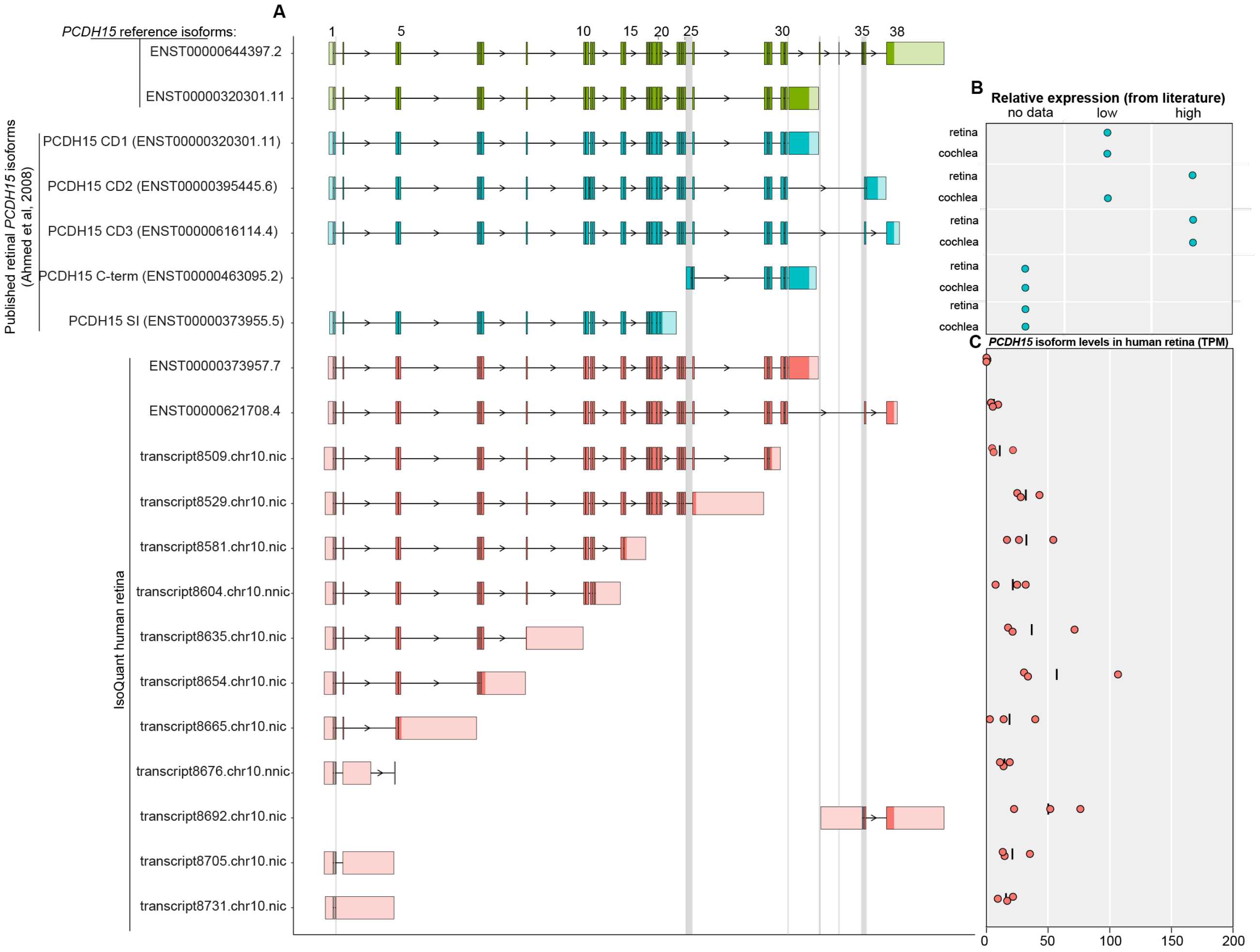


**Figure S5: *PCDH15* isoforms identified by IsoQuant analysis compared to known isoforms from the literature. A.** The GENCODE reference transcripts are depicted at the top in green, followed by the known human *PCDH* isoforms in blue (Ahmed et al., 2008). The *PCDH15* IsoQuant transcripts are depicted in red. The light green, blue and red colors indicate the untranslated regions (UTR) and the dark green, blue and red colors indicate the open reading frame (ORF) of each transcript. Differences between the IsoQuant isoforms and the GENCODE reference transcript are highlighted in grey boxes. **B.** Relative expression of *PCDH15* isoforms based on literature in either the retina or the cochlea. **C.** The Transcripts Per Million (based on dataset 1) for the IsoQuant identified isoforms are presented for the three individual samples.

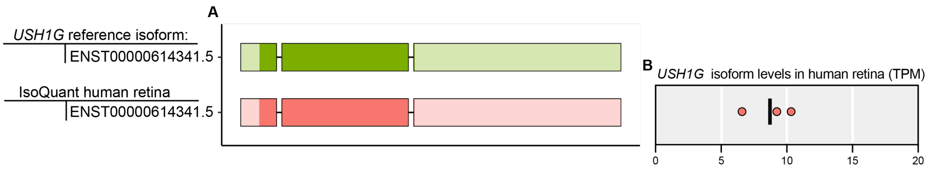


**Figure S6: *SANS* isoform identified by IsoQuant analysis compared to known isoform. A.** The GENCODE reference transcript is depicted at the top in green, followed by the *SANS* IsoQuant transcript depicted in red. The light green and red colors indicate the untranslated regions (UTR) and the dark green and red colors indicate the open reading frame (ORF) of each transcript. **B.** The Transcripts Per Million (based on dataset 1) for the IsoQuant identified isoform are presented for the three individual samples.

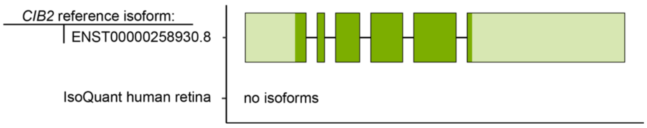


**Figure S7: No IsoQuant isoforms are identified for *CIB2*** The GENCODE reference transcript is depicted at the top in green with light green colors indicating the untranslated regions (UTR) and the dark green colors indicating the open reading frame (ORF) the reference isoform. No IsoQuant isoforms are identified for *CIB2.*

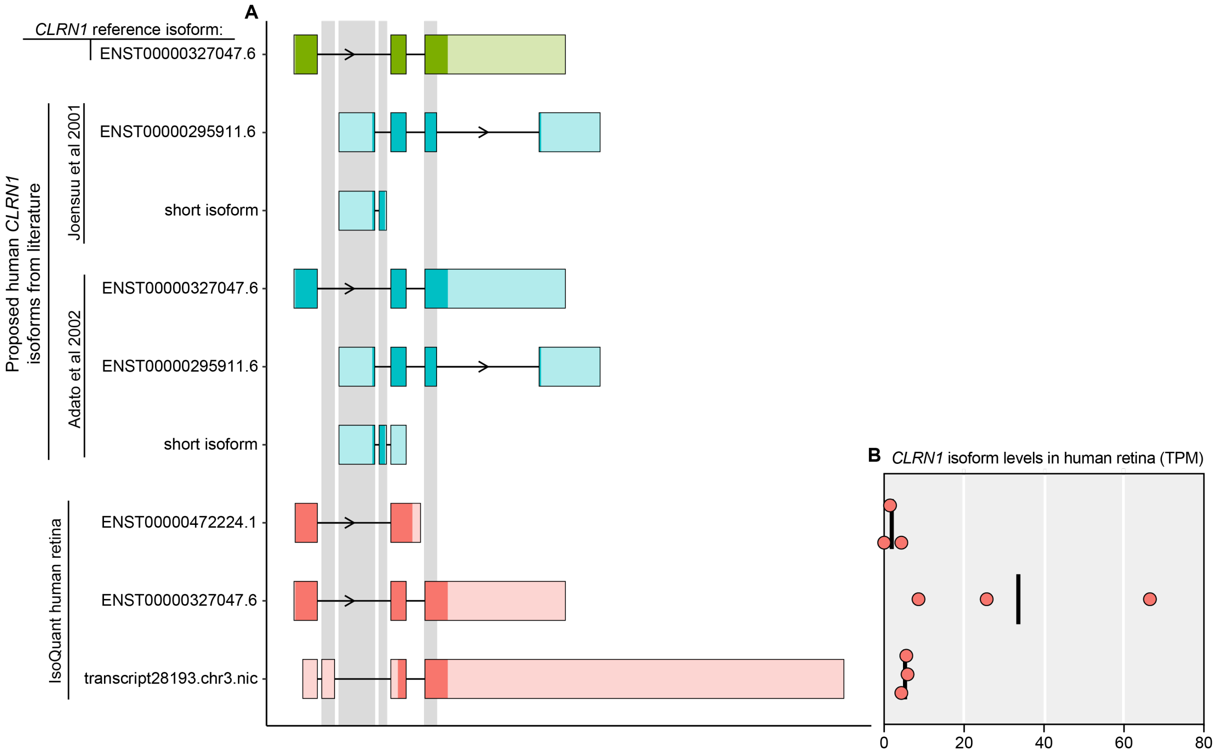


**Figure S8: *CLRN1* isoforms identified by IsoQuant analysis compared to known isoforms from the literature. A.** The GENCODE reference transcript is depicted at the top in green, followed by the known human *CLRN1* isoforms in blue (Joensuu et al., 2001 Adato et al., 2002). The *CLRN1* IsoQuant transcripts are depicted in red. The light green, blue and red colors indicate the untranslated regions (UTR) and the dark green, blue and red colors indicate the open reading frame (ORF) of each transcript. Differences between the IsoQuant isoforms and the GENCODE reference transcript are highlighted in grey boxes. **B.** The Transcripts Per Million (based on dataset 1) for the IsoQuant identified isoforms are presented for the three individual samples.

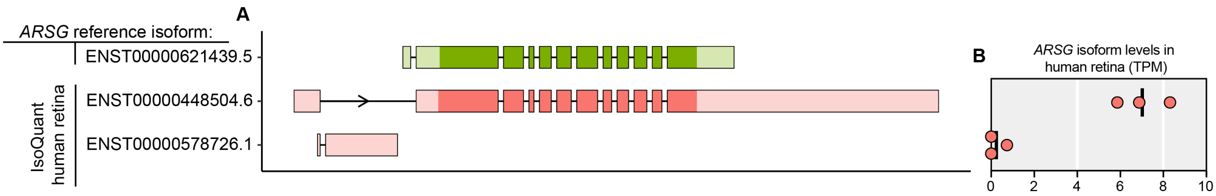


**Figure S9: *ARSG* isoforms identified by IsoQuant analysis compared to known isoform. A.** The GENCODE reference transcript is depicted at the top in green, followed by the *ARSG* IsoQuant transcripts depicted in red. The light green and red colors indicate the untranslated regions (UTR) and the dark green and red colors indicate the open reading frame (ORF) of each transcript. **B.** The Transcripts Per Million (based on dataset 1) for the IsoQuant identified isoforms are presented for the three individual samples.
